# Neural mechanisms to incorporate visual counterevidence in self motion estimation

**DOI:** 10.1101/2023.01.04.522814

**Authors:** Ryosuke Tanaka, Baohua Zhou, Margarida Agrochao, Bara A. Badwan, Braedyn Au, Natalia C. B. Matos, Damon A. Clark

**Author notes:** Institute of Neuroscience, Technical University of Munich, Munich 80802, Germany (Ryosuke Tanaka).

## Abstract

In selecting appropriate behaviors, animals should weigh sensory evidence both for and against specific beliefs about the world. For instance, animals measure optic flow to estimate and control their own rotation. However, existing models of flow detection can confuse the movement of external objects with genuine self motion. Here, we show that stationary patterns on the retina, which constitute negative evidence against self rotation, are used by the fruit fly *Drosophila* to suppress inappropriate stabilizing rotational behavior. *In silico* experiments show that artificial neural networks optimized to distinguish self and world motion similarly detect stationarity and incorporate negative evidence. Employing neural measurements and genetic manipulations, we identified components of the circuitry for stationary pattern detection, which runs parallel to the fly’s motion- and optic flow-detectors. Our results exemplify how the compact brain of the fly incorporates negative evidence to improve heading stability, exploiting geometrical constraints of the visual world.

## Introduction

Animals constantly monitor their sensory inputs to detect significant events so that they can respond with appropriate actions. However, sensory inputs are often open to multiple interpretations. When evaluating the likelihood of a certain event occurring given sensory inputs, it is important to incorporate not only the evidence *confirming* such interpretation, but also evidence *against* the interpretation. Failure to acknowledge negative evidence is well documented in the domain of human reasoning^1^ as well as in perceptual decision making tasks^2, 3^, a phenomenon termed confirmation bias. However, for all animals, initiating spurious behavioral responses to incorrectly interpreted sensory inputs could be very costly. Therefore, one might expect that animals be able to weigh negative evidence better in contexts of innate, naturalistic behaviors.

A critical innate behavior that is based on visual inferences is course stabilization. When an animal moves relative to its environment, it experiences systematic, panoramic patterns of visual motion, called optic flow^4^. Diverse species, ranging from vertebrates to invertebrates, estimate their own movement based on optic flow and use that information to stabilize their locomotion^5, 6^. Because animals need to enact their intended movements if they are to achieve their behavioral goals, the algorithms they use to detect visual motion and estimate their own velocity are often regarded as highly optimized for accuracy^7^. Reflecting the importance of the optic flow-based course stabilization behaviors, neurons tuned to specific patterns of optic flow have been found in the brains of diverse species across taxa^8–12^. These flow sensitive neurons are thought to achieve their selectivity by weighting local motion cues over space to match them with template flows generated by different types of self motion, an algorithm termed template matching^13, 14^. Template matching can effectively distinguish different types of optic flow, including by negatively weighting local visual motion oriented against the flow template, a process whose microcircuit implementation has been well studied in insects^15, 16^.

However, these template matching algorithms can overlook the alternative possibility that the observed motion did not result from the movements of the observer. For example, a large object traversing in front of a stationary observer will generate a pattern of visual motion largely consistent with optic flow during yaw rotation (**Figure 1a**). This sort of “world motion” scene will spuriously trigger template matching-based yaw detectors, since it does not contain local visual motion inconsistent with the yaw optic flow template. Incorrectly initiating stabilizing behaviors in response to world motion will deviate animals from their intended courses and is thus maladaptive. We therefore hypothesized that animals have evolved additional strategies to distinguish visual motion caused by world motion from genuine optic flow. An important visual cue to distinguish between self rotation and world motion is patterns that are stationary on the retina. The geometry of optic flow during yaw rotation dictates that visual motion is oriented in the same direction everywhere and that the speed is constant across all points sharing an elevation. Therefore, the existence of any stationary pattern in the visual field strongly implies that the observer is not, in fact, rotating (**Figure 1b**), even when parts of the visual field are moving coherently. Importantly, motion-based template matching algorithms are insensitive to this form of negative evidence against self motion, because areas with zero velocity do not contribute either positively or negatively to the total weighting of the visual motion field.

**Figure 1.**
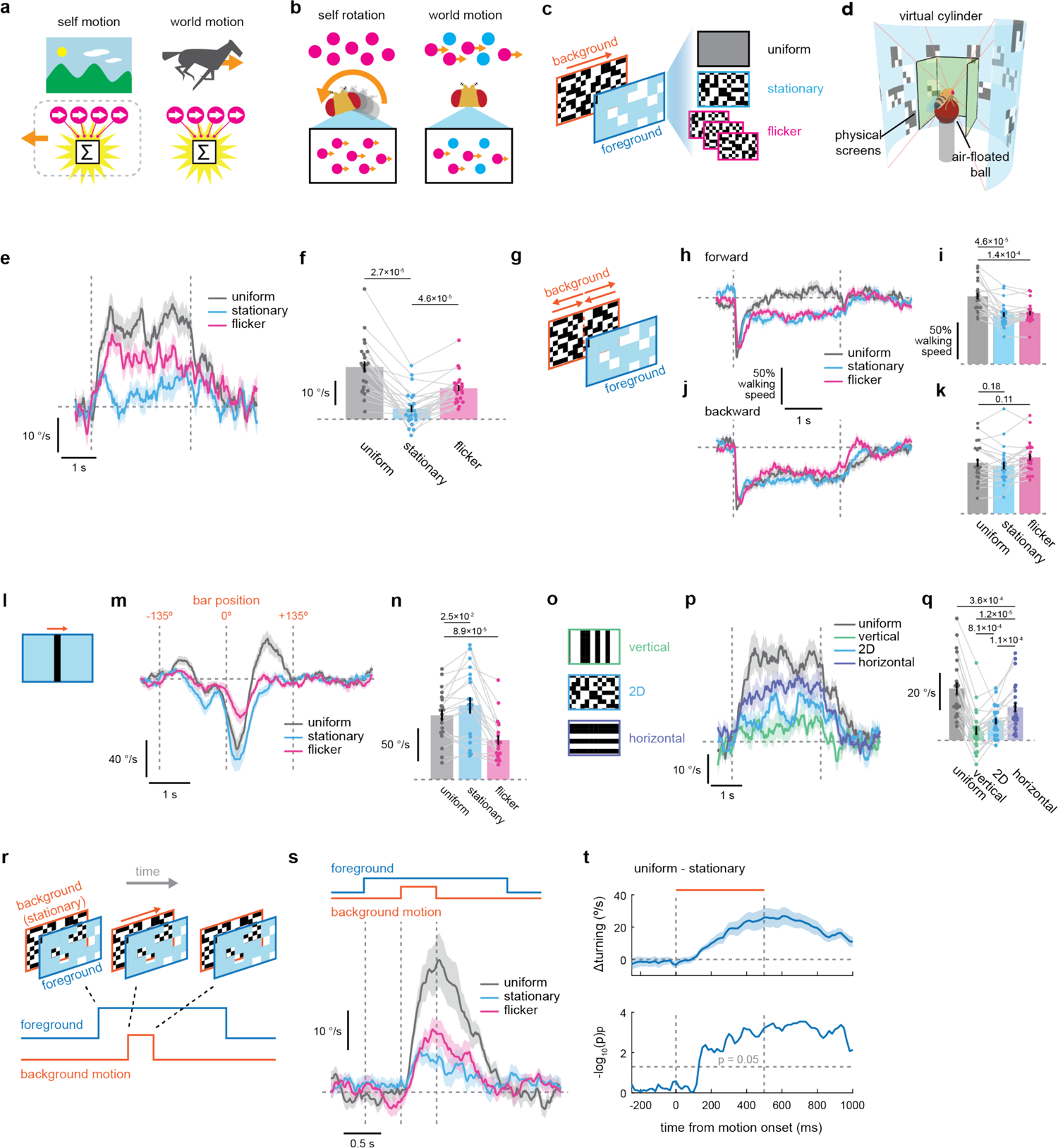
*Drosophila* use stationary patterns as negative evidence against self rotation to suppress rotational optomotor responses. (a) Self motion and world motion can both activate a template matching-based flow detector in a similar way. **(b)** When an observer rotates in the environment, every visible object moves in the same direction. Therefore, the existence of any stationary patterns in the visual field argues against self rotation. **(c)** A schematic of the islands of motion stimuli. Rotating background was viewed through windows in the foreground. **(d)** A schematic of the behavioral setup. Flies experienced visual stimuli on a virtual cylinder through a projection onto rectangular panoramic screens surrounding them. **(e, f)** Wild type fly turning response to the islands of motion stimuli (e) over time and (f) averaged over time. Vertical lines mark the onset and offset of the stimuli, and the horizontal line marks zero turning. Throughout, positive angular velocities indicate turning in the same direction as the stimulus. Individual dots in (f) represent individual flies, and data from the same flies are connected with gray lines. **(g)** In the translational islands of motion stimuli, the background moves front-to-back or back-to-front in a symmetric fashion about the fly. **(h-k)** Normalized walking speed of flies in response to the (h, i) forward or (j, k) backward translational islands of motion stimuli, either over time (h, j) or averaged over time (i, k). Horizontal dotted lines indicate 100% walking speed during the pre-stimulus period. Data in (h-k) are from the same set of flies as in (e, f). **(l)** A schematic of the moving bar stimulus. The blue “foreground” region was filled with the same three foreground patterns (uniform, stationary, or flicker) as in (c) and (g). **(m, n)** Turning responses of flies in response to the bar stimuli, either (m) over time or (n) peak response amplitude. **(o)** Schematic of stimuli used to probe orientation selectivity of the optomotor suppression caused by stationary foreground patterns. **(p, q)** Fly turning responses to islands of motion stimuli with one-dimensional stationary foreground patterns, either (p) over time or (q) averaged over time. **(r)** A schematic of the modified islands of motion stimuli. The background and foreground patterns appear simultaneously, but the motion starts after a brief delay. **(s)** Fly turning response to the modified islands of motion stimuli in (r) over time. Vertical lines mark the onset of stimulus, the onset of motion, and the offset of motion. The horizontal line marks zero turning. The onset of background motion was delayed 0.5 s and background motion lasted 0.5 s. **(t)** (top) Difference of fly turning responses to islands of motion stimuli with uniform and stationary foreground patterns, averaged across flies. Vertical dotted lines mark the onset and offset of visual motion. The orange horizontal line marks the period during which the background was moving. (bottom) Negative of log10 transformed Wilcoxon signed-rank p-values between the uniform and stationary foreground conditions across flies, computed at each time point. The horizontal dotted line mark p = 0.05. Shading around the time traces and error bars on the bar plots indicate standard error of the mean across flies. (e-k) N = 23 flies. (m, n) N = 20 flies. (p, q) N = 23 flies. (s, t) N = 17 flies. Numbers over bar plots indicate p-values from two-sided Wilcoxon signed-rank tests.

In the present study, we have discovered that the fruit fly *Drosophila* uses stationary visual patterns as negative evidence against self rotation, as predicted from the geometrical argument above. With high-throughput behavioral assays, we find that stationary visual patterns selectively suppress the rotational course stabilization behaviors of the flies, but not other types of motion-dependent and turning behaviors. In a parallel *in silico* experiment, we show that simple artificial neural networks optimized to distinguish between genuine optic flow and object motion generate similar behavioral patterns to those observed in flies. By performing two-photon calcium imaging on the identified visual circuits for motion and optic flow detection, we demonstrate that integration of evidence for and against self rotation takes place in the central brain, downstream of flow-sensitive peripheral neurons, which are largely unaffected by stationary patterns. Finally, through a behavioral genetic screen, we identified genetically defined populations of neurons necessary for the suppression of optomotor response in the presence of stationary patterns, which include the early visual neuron Mi4. A simple model based on Mi4 response properties can achieve sensitivity to stationary patterns. Overall, our results demonstrate how flies exploit the geometrical constraints of the visual world to incorporate both positive evidence for and negative evidence against self rotation as they stabilize their orientation in the world.

## Results

### Drosophila suppress rotational optomotor responses in the presence of stationary patterns

When flies are presented with wide-field visual patterns rotating around them in the yaw direction, they exhibit stereotyped turning responses in the same directions as the visual motion, a behavior termed the rotational optomotor response^17, 18^. To test whether flies utilize stationary patterns as negative evidence against self rotation to suppress optomotor response, we designed stimuli we called “islands of motion” (**Figure 1c, Supplementary Video 1**). In these stimuli, a background of a random binary checkerboard with 5° resolution rotated in yaw at 80°/s around the observer and was paired with different foreground patterns. Foreground patterns had transparent square-shaped windows through which the moving background was visible, creating “islands” of visual motion. The windows were randomly placed, had the size of 15°, and covered an average of 20% of the visual field. The foreground pattern was either (1) uniform gray (labeled ‘uniform’), (2) a stationary 5° random binary checkerboard (labeled ‘stationary’), or (3) a 5° random binary checkerboard randomly updated at 15 Hz (labeled ‘flicker’). The sizes of checkerboard patterns (5°) and windows (15°) were respectively chosen to be close to the receptive field sizes of photoreceptors^19^ and elementary motion detectors^20, 21^ in *Drosophila*. Importantly, because the foreground patterns contained no net directional motion and the relative area of the windows were kept constant across the three conditions, the stimuli contained the identical total directional motion regardless of the foreground pattern. However, only the stationary foreground condition contains stationary patterns that can be potentially used as negative evidence against self rotation.

To measure fly optomotor responses to the islands of motion stimuli, we tethered wild-type flies on surgical needles and allowed them to walk on air floated balls. Flies viewed the visual stimuli presented on a simulated cylindrical wall surrounding them through projections onto small planar screens; the appropriate image transformations were achieved with OpenGL (**Figure 1d**)^22^. We discovered that the existence of a stationary pattern in the foreground resulted in almost 10-fold suppression of the optomotor response amplitude relative to the uniform foreground condition (**Figure 1e, f**). This is consistent with the hypothesis that flies integrate negative visual evidence into rotational optomotor behavior. Notably, the suppression of turning by stationary patterns was even stronger than the suppression caused by flickering patterns, which are known to suppress motion detector activity, and thus optomotor turning, through a spatial contrast normalization mechanism^23^.

### Stationary patterns specifically suppress optomotor turning responses

The observed suppression of optomotor response is consistent with the functional interpretation that flies are specifically using stationary patterns as negative evidence against self rotation. However, it is also possible that stationary patterns are suppressing all motion-dependent behaviors, not just responses to rotational optic flow. To test this possibility, we asked if stationary patterns also suppress other motion-dependent behaviors and turning behaviors. First, we focused on the visual control of forward walking speed. Walking flies accelerate or decelerate in response to translational optic flow to stabilize their walking speed, a behavior that depends on the same early motion detecting neurons as the optomotor turning response^24, 25^. In an open-loop setting, forward and backward optic flow both make flies decelerate^24, 25^. To probe if the walking speed control is also similarly suppressed by stationary patterns, we paired the same set of foreground patterns with background patterns moving symmetrically about the fly, which approximated translational optic flow during forward or backward translational movements (**Figure 1g**). Importantly, unlike in the case of self rotation, stationary patterns do not imply the lack of self translation. This is because far away objects can remain visually stationary during translation. Thus, there is no geometrical reason to expect stationary patterns to suppress forward walking speed modulation in response to translational optic flow. Here, we observed that both stationary and flickering foregrounds increased, rather than decreased, the strength of the slowing caused by forward optic flow (**Figure 1h, i**). They did not significantly affect slowing caused by backward optic flow (**Figure 1j, k**).

Next, we asked if the stationary foreground suppresses flies’ visually driven turning in response to moving visual objects. When presented with isolated moving vertical bars, flies turn in the same^26–29^ or opposite direction^30, 31^, depending on the speed of the bar, a behavior that depends on the same set of early motion detecting neurons as the optomotor turning response^28, 30, 31^. Unless stationary patterns broadly suppress the motion detectors or turning, there is no geometrical reason to expect stationary patterns to suppress turning in response to moving objects. To test this idea, we presented a fast-moving vertical bar on the same three (uniform, stationary, or flicker) foreground patterns (**Figure 1l**). The peak turning amplitude was not decreased by stationary patterns, while flickering patterns suppressed the turning (**Figure 1m, n**), consistent with spatial contrast normalization in motion detectors^23^. Overall, these results show that the suppression of turning by stationary patterns is specific to optomotor response, supporting the functional interpretation that flies are using the stationary patterns as negative evidence against self rotation.

### The suppression of rotational optomotor responses mirrors the geometry of self rotation

If flies are using stationary patterns as negative evidence against self rotation, we can make further predictions about what kinds of foreground patterns should mostly strongly suppress rotational optomotor responses. During yaw rotation, vertical edges move sideways, while horizontal edges appear to remain stationary on the retina. As a consequence, whereas stationary vertical patterns strongly argue against self rotation, stationary horizontal patterns are uninformative about whether or not the observer is rotating. To test whether the suppression of optomotor responses shows the pattern of orientation selectivity expected from this geometrical rule, we presented islands of motion stimuli with stationary foregrounds in patterns that were either vertical, two-dimensional, or horizontal (**Figure 1o**). As predicted by the geometrical reasoning above, vertical stationary patterns suppressed turning more than two-dimensional or horizontal patterns, further buttressing the functional interpretation that flies are using stationary patterns as negative evidence against self rotation (**Figure 1p, q**). Flickering horizontal and vertical patterns also resulted in similar orientation selective suppression of optomotor response (**Figure S1a-c**).

So far, we have focused on the use of contrast-defined stationary patterns as negative evidence against self rotation. Another geometrical cue that signifies an absence of self motion is stationary motion-defined contours, that is, boundaries between areas with different velocities that are themselves stationary. During genuine self rotation, regions of the visual field containing visual motion should rotate their position consistently with visual motion inside them. That is, the “islands of motion” should appear to be sliding over space (**Figure S1d**). Conversely, the existence of stationary islands of motion in our stimuli argues against self rotation, even when the foreground does not contain contrast-defined stationary patterns. We think of this cue as second-order^32^, since it concerns spatiotemporal correlations in a higher-order image statistic, namely local motion. Flies have been shown to be able to detect certain forms of second-order motion when tracking objects in flight^33^. To test whether flies can use this second-order cue for the absence of self rotation, we presented flies with islands of motion stimuli where the islands either remained fixed or moved with the rotation of the background (**Figure S1d**). Flies turned slightly but significantly more to the stimuli with sliding windows than to the ones with fixed windows (**Figure S1e, f**). In comparison, a contrast-defined stationary foreground pattern strongly suppressed turning even when the windows were sliding (**Figure S1e, f**). This observation suggests that flies do not use motion-defined stationary contours as strong negative evidence against self rotation, especially compared to contrast-defined stationary patterns.

### Optomotor suppression is as fast as optomotor response itself

Until this point, we have characterized the suppression of optomotor response in terms of the time averaged turning amplitude over several seconds. While it is important that the suppression can last for many seconds in the presence of lasting world motion, it is also critical for course stability that the suppression can act fast. In the initial behavioral experiment (**Fig. 1e**), the suppression becomes apparent more than 300 ms after the stimulus onset, which is slower than the rise time of optomotor response itself (**Fig. S1g, h**). However, in a more naturalistic setting, stationary patterns do not suddenly appear simultaneously with visual motion, as in our initial stimulus design (**Supplementary Video 1**). Rather, in natural settings, some stationary visual patterns would tend to be visible before world motion occurs, a situation that could make it possible for flies to suppress spurious optomotor response faster than we observed in our initial experiments (**Fig. S1g**). To test this idea directly, we redesigned our islands of motion stimuli so that the background motion starts only after a brief delay (**Fig. 1r**). In this setting, turning responses to uniform and stationary conditions started to diverge earlier, reaching statistical significance ∼150 ms after the motion onset (**Figs. 1s, t, S1i, j**). This time course is comparable to the observed rise kinetics of optomotor turning itself, indicating that the optomotor suppression can act fast.

### Artificial networks trained to distinguish self and world motion exhibit behavior similar to flies

Based on geometrical arguments, we have so far interpreted the observed suppression of optomotor response by stationary visual patterns as a manifestation of a strategy to prevent inappropriate optomotor response in the face of world motion. However, such functional, adaptationist claims are in principle difficult to prove, and require careful consideration^34^. In order to better interpret the fly behavior, we decided to compare it to the behavior of artificial agents trained to distinguish self and world motion using modern machine learning techniques^35^. If such trained artificial neural networks (ANNs) develop a similar strategy to use stationary visual patterns as negative evidence against self motion, that lends additional credence to our functional interpretation of the fly behavior. In addition, by analyzing the trained ANNs, we might gain insight into how an algorithm to detect stationary patterns could be implemented.

Here, we designed ANNs with varying numbers of layers and channels, and trained them to distinguish videos of self and world motion (**Figure 2a**). In a biological analogy, each channel corresponds to a local, columnar visual neuron cell type with a unique, spatially-invariant receptive field structure, and each of the *L* layers of an ANN housed *C* different cell types. The layer and channel numbers *L* and *C* were respectively varied in the range of 1 to 6 and 1 to 8. Importantly, the ANNs had a convolutional architecture, where each virtual cell type tiled the entire visual space with a spatially invariant receptive field. The training samples provided to the ANNs were videos of naturalistic scenes representing either self or world motion. The videos were created by extracting single horizontal slices from monochromatic, panoramic images of natural scenes^36^ (**Figure 2b**). In the self motion videos, the whole scene translated rigidly in the horizontal direction with time-varying velocity drawn from an autocorrelated Gaussian process^37, 38^. In the world motion videos, random patches of contrast taken from different regions of a natural scene mimicked external objects that moved horizontally across the stationary background scene. The velocities of the patches were drawn from the same process as the self motion videos. The ANNs were then trained to estimate the probability that the video belonged to the self motion class (see *Materials and Methods* for details). For each architecture (*i.e.,* each *L* and *C* combination), we trained 100 instances of models with random initialization. Here, each of the 100 model instances can be envisioned as different “species” with the identical architectural constraint on their visual system.

**Figure 2.**
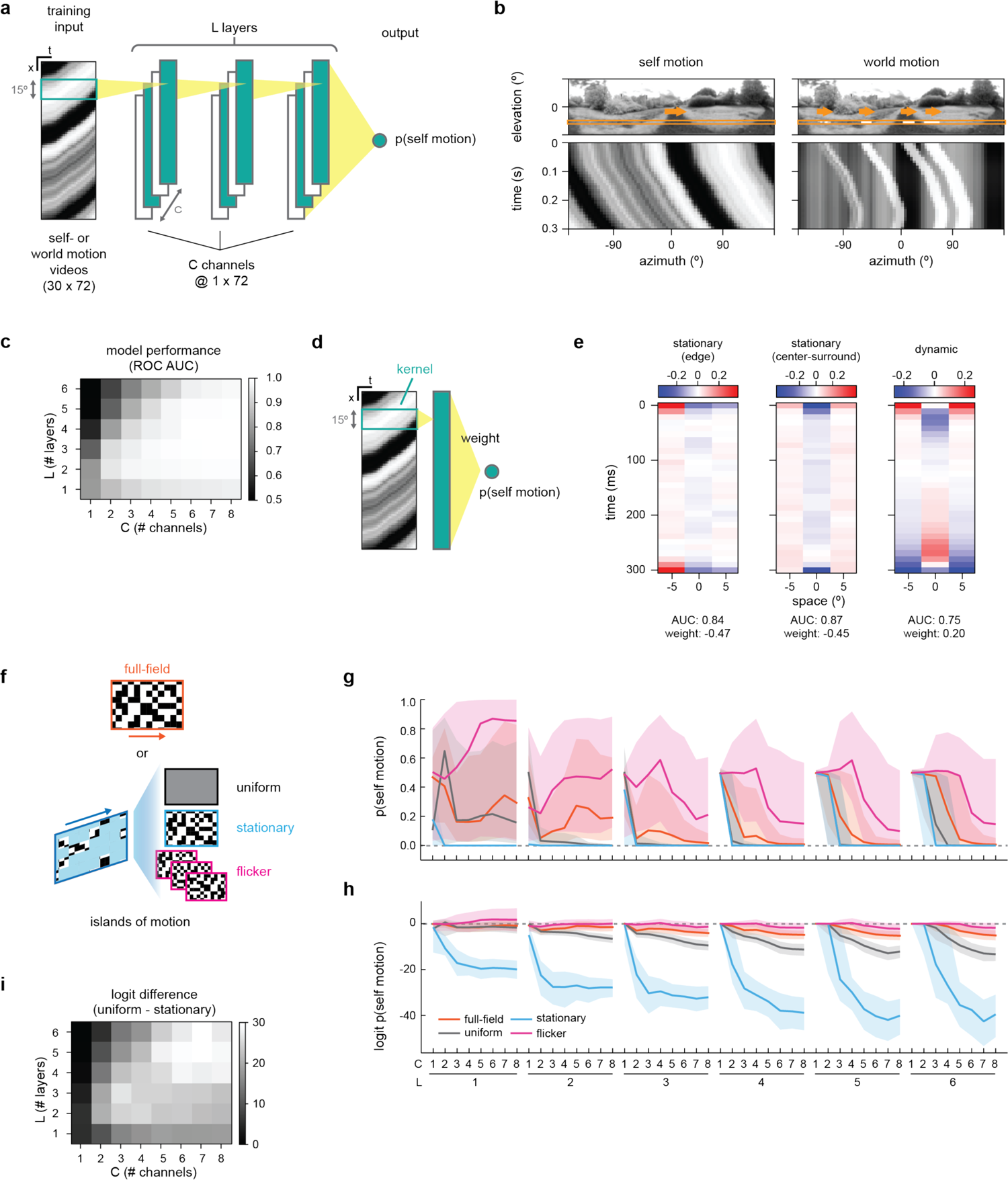
A simple artificial neural network trained to distinguish self and world motion exhibits similar behaviors to flies. **(a)** A schematic of the model architecture and training. The model consisted of *L* convolutional layers with *C* channels. The spatial width of the convolutional kernels was fixed at 15°, three pixels with 5° spacing. The depth of the kernel was the duration of the video (30 frames) for the first layer, and the number of channels in the following layers. The activity of the last convolutional layer was averaged with spatially uniform weights and logistic-transformed to generate the estimate of the probability that the input video belongs to the self motion condition. **(b)** Illustrative examples of videos used for training artificial neural networks (ANNs). The videos were created by taking out single horizontal slices of panoramic images of natural scenes. For the self motion condition, the entire scene moved horizontally following a stochastic velocity time series. In the world motion condition, only “objects” moved and the scene itself remained stationary, where the objects are patches randomly selected from the same image. A single world motion image contained up to 10 objects, with widths of each between 5° to 36°. **(c)** The average model performance for each model architecture quantified as the area under the curve (AUC) of the receiver-operator characteristic (ROC) curve, calculated on the held-out test dataset. **(d)** A schematic of the simplest model with *L* = 1 and *C* = 1. **(e)** Representative examples of the spatiotemporal kernels from the trained models with the simplest architecture, as shown in (d). AUC scores corresponding to the kernels are below the heatmaps. The signs of the weights indicate how the kernels contributed to the estimated probability of self motion. See also **Figure S2a-c**. **(f)** A schematic of the simulated experiment. Trained models were presented with the full-field moving checkerboard or the islands of motion stimuli as in **Figure 1c**. **(g)** The predicted probability of self motion for the full-field rotating checkerboard or the islands of motion stimuli, for different model architectures. The lines represent median, and shaded regions represent 25^th^ and 75^th^ percentile performances across 100 model initializations. **(h)** Same as (g), but showing the logit values instead of the probability of self motion. **(i)** Difference between the logit values of the self motion probability between the uniform and the stationary conditions, shown as functions of the model architecture.

The trained ANNs on average distinguished well between self and world motion, as quantified with the area under the curve (AUC) metrics of the receiver-operator characteristic (ROC) curves^39^ (**Figure 2c**). The exception was the deepest models with small numbers of channels (*e.g.*, *L* = 6, *C* = 1). The poor performance of these networks represents the failure of training rather than the intrinsic limitation of its network architecture, because they can in principle implement identical functions as shallower, better-performing networks. In fact, the simplest network with only a single layer and a single channel was able to perform the task with an AUC of approximately 0.8.

To better understand the algorithms used by the trained networks to distinguish between self and world motion, we visualized the convolutional kernels of some of the top-performing instances of the model with the simplest architecture (i.e. *L* = 1, *C* = 1) (**Figure 2d, e, S2a-c**). The kernels of the first layer of the networks are directly convolved with the input images, and thus can be straightforwardly interpreted as linear receptive fields. Since there were no biophysical constraints in the model, the filters tended to spread their weight to the two ends of the filter. Nevertheless, the filters remain interpretable by their properties, as we discuss below. Out of the 100 randomly initialized instances of the model (*i.e.,* “species”), roughly a half (46 of 100) had kernels with temporally constant polarities (static kernels), whereas the rest (54 of 100) had kernels with changing polarity (dynamic kernels) (**Figure 2e, S2a-c**). The static kernels had an edge-detector or center-surround antagonistic structure, making them sensitive to stationary patterns. Their outputs contributed negatively to the estimated probability of self motion (**Figure 2e, S2a-c**). Thus, this first class of models with static kernels used stationary patterns as negative evidence against self motion, similar to flies. In contrast, the dynamic kernels had a center-surround structure with preferred contrasts that flipped between the first and second halves of the kernel (**Figure 2e, S2a-c**), making them sensitive to changes in contrast over time, which is associated with scene motion. The dynamic kernels contributed positively to the probability of self motion. Thus, the models with dynamic kernels use contrast changes as positive evidence in favor of self motion. Importantly, the models with stationary kernels always substantially outperformed ones with dynamic kernels (stationary: AUC = 0.857 ± 0.012, dynamic: AUC = 0.755 ± 0.007, mean ± standard deviation).

Next, we tested how the trained ANNs respond to the islands of motion stimuli used in the behavioral experiments, as well as to full-field rotating checkerboards (**Figure 2f**). To this end, single horizontal slices of these stimuli were downsampled to the 5° and 100 Hz spatiotemporal resolution of the ANN simulations. In general, successfully trained ANNs predicted the highest probability of self motion for the full-field rotating checkerboards as well as for the islands of motion stimuli with flickering foregrounds (**Figure 2g, h**). Most importantly, the trained ANNs predicted lower probability of self motion for the islands of motion stimuli with stationary foregrounds compared to ones with uniform foregrounds, consistent with the behavior of flies. The difference of responses between the uniform and stationary conditions, as quantified by logit differences (**Figure 2i**), increased with the number of channels. Upon closer examination of the simplest (*L* = 1, *C* = 1) models, we found that only models with stationary kernels, but not ones with dynamic kernels, were able to predict low probability of self motion in the presence of stationary patterns, highlighting the difference between the positive and negative evidence detection strategies (**Fig S2d**). Unlike flies, the trained ANNs strongly distinguished between the islands of motion stimuli with fixed and sliding windows (**Figures S1d-f, S2e, f**). Overall, these observations support the idea that systems optimized to distinguish self and world motion in natural scenes can acquire a strategy to use stationary patterns as negative evidence against self motion. In particular, simple yet successful models used spatially derivative-taking and temporally sustained kernels to detect stationary patterns, hinting at how equivalent computations might be implemented in the fly brain.

### Characterizing the visual tuning of optomotor suppression

Next, in order to confirm that the behavior under study is robust across a variety of stimulus parameters as well as to constrain underlying neural mechanisms, we performed additional psychophysical experiments on wild-type flies to characterize how the observed suppression of optomotor response depended on different stimulus parameters. First, we swept the contrast of foreground patterns. As naively expected, the amount of suppression decreased as the contrast decreased, both for stationary and flickering patterns (**Figure 3a**). Second, we modulated the fraction of the visual scene occupied by background and foreground. We observed significant suppression of turning for both flickering and stationary foreground types when the islands of motion covered up to 40% of the screen (**Figure 3b**). Third, we found that suppression of turning by both flickering and stationary foreground patterns did not depend strongly on the island velocity (**Figure 3c**). Fourth, we modulated the update rate of the flickering foreground, creating a continuum between the stationary and flickering conditions. We found that the optomotor suppression peaked at an update rate of ∼5 Hz (**Figure 3d**). Fifth, to probe the spatial frequency tuning of a putative stationary pattern detector, we used stationary plaid patterns generated by superimposing vertical and horizontal sinusoidal gratings with various wavelengths as the foreground pattern. The amount of suppression increased until the wavelength of 30° and then stayed high up to about 60° (**Figure 3e**), revealing a band-pass property. Sixth, we modulated the mean luminance of the foreground patterns to probe the contrast preference of the hypothetical stationary pattern detectors. We found that half-contrast stationary foreground with luminance below the mean background luminance recapitulated the full extent of turning suppression caused by the full-contrast foreground (**Figure 3f**). This cannot be explained as the simple effect of overall luminance decrement, because uniform darker foreground increased the optomotor response, rather than decreasing it, relative to the uniform mean-gray foreground (**Figure 3g**). This result suggests that the hypothetical stationary pattern detector is more sensitive to local contrast decrements than increments.

**Figure 3.**
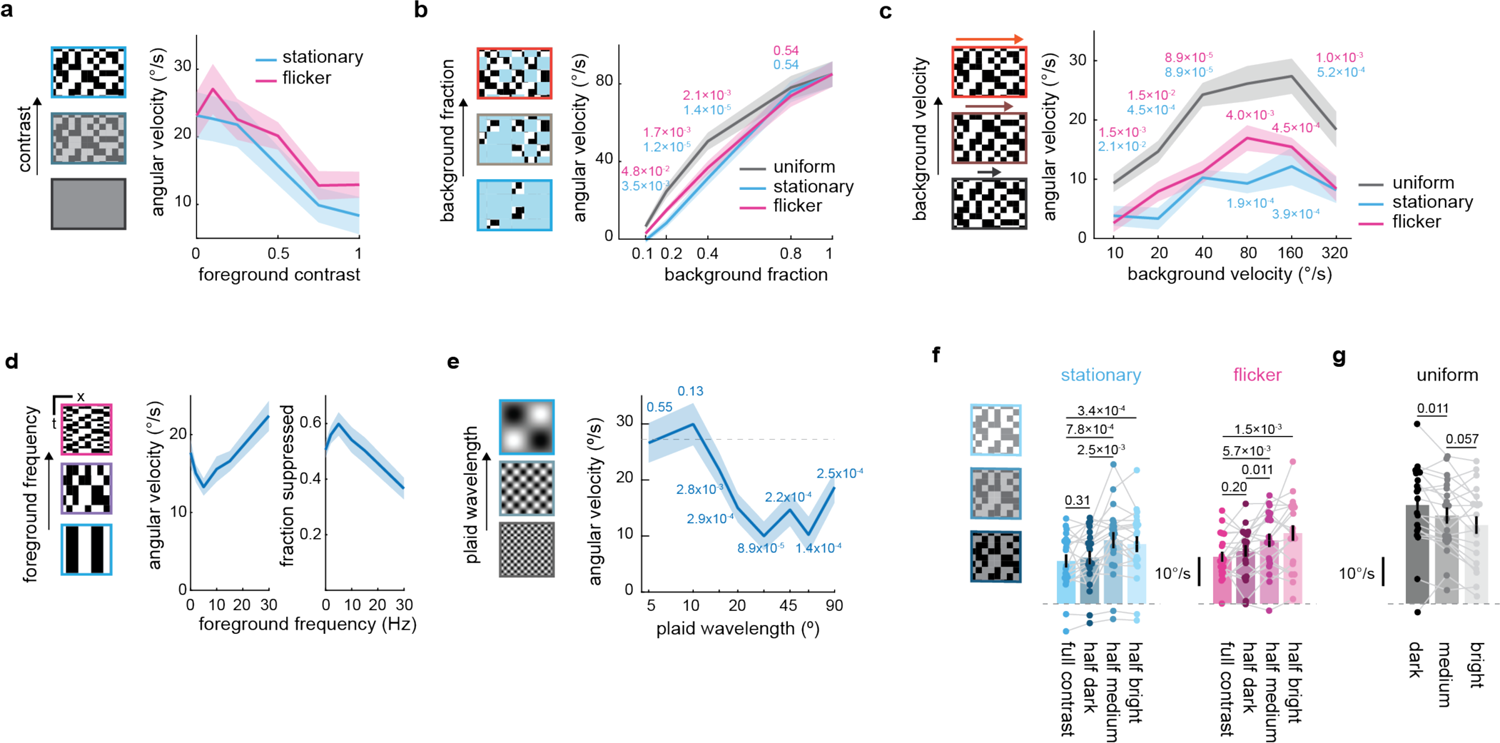
Visual tuning of the optomotor suppression by stationary patterns. **(a)** Time-averaged turning responses of flies to the islands of motion stimuli with stationary or flickering foreground patterns with varying foreground contrasts. **(b)** Time-averaged turning responses of flies to the islands of motion stimuli with different fractions of background coverage. **(c)** Time-averaged turning responses of flies to the islands of motion stimuli with different background velocities. **(d)** Time-averaged turning responses of flies to the islands of motion stimuli with flickering foreground patterns updated at different temporal frequencies. (*left*) Raw turning amplitude, and (*right*) fractional suppression relative to the uniform foreground condition. **(e)** Time-averaged turning responses of flies to the islands of motion stimuli with plaid foreground patterns with different spatial wavelengths. The horizontal dotted line indicates the turning amplitude in the uniform foreground condition, and numbers indicate p-values from signed-rank tests between the uniform and different plaid conditions. **(f)** Time-averaged turning responses of flies to the islands of motion stimuli with full- and half-contrast foreground patterns with different mean luminance. **(g)** Time-averaged turning responses of flies to the islands of motion stimuli with the uniform foregrounds with different luminance levels. (a) N = 19 flies. (b) N = 25 flies. (c) N = 20 flies. (d) N = 19 flies. (e) N = 20 flies. (f, g) N = 20 flies. Numbers over plots indicate p-values from two-sided Wilcoxon (b, c) rank-sum or (e-g) signed-rank tests. Cyan and magenta numbers in (b, c) respectively indicate the results of statistical tests between uniform vs. stationary or uniform vs. flicker conditions.

### Stationary patterns do not suppress elementary motion detectors

We next sought to identify the neural bases of the optomotor suppression by stationary patterns. A straightforward hypothesis is that there are dedicated visual neuron types that detect stationary patterns and subsequently suppress neurons somewhere in the neural pathway implementing the optomotor response (**Figure 4a**). To investigate this possibility, we first used two-photon calcium imaging to attempt to find neurons in the optomotor response pathway that are suppressed by stationary patterns (**Figure 4b**). To start, we recorded the calcium activity of T4 and T5 neurons. T4 and T5 neurons are the first direction selective neurons in the fly visual system and are necessary for rotational optomotor responses^40^ (**Figure 4c**). To record the activity of individual T4 and T5 axon terminals in lobula plate, we sparsely expressed a genetically encoded calcium indicator GCaMP6f^41^ in T4/T5 neurons using SPARC, a genetic method to achieve stochastic transgene expression under the Gal4 control^42^. As visual stimuli, we used a modified version of the islands of motion stimuli, where the background was presented in a single circular aperture with 15° diameter, centered about the receptive field of individual T4 and T5 cells (**Figure 4d**). We set a 5° wide uniform gray buffer zone between the aperture and the foreground to prevent foreground patterns from overlapping with the receptive fields of T4 and T5. We found that stationary foreground patterns significantly suppressed T5 responses by about 25%, whereas T4 responses were not affected by the stationary patterns, for both preferred (**Figure 4e, f**) and non-preferred directions (**Figure S3a, b**). In contrast, flickering foreground patterns strongly suppressed the activity of both T4 and T5 (**Figure 4e, f**). The observation that dynamic patterns in the surround suppress T4 and T5 activities is consistent with spatial contrast normalization^23^. Overall, while stationary patterns did slightly suppress T5 activities, the amplitude of the suppression appears too small to explain the behavioral observations (i.e., 90% suppression of turning) (**Figure 1**). Importantly, this observation is consistent with the behavioral finding that motion-dependent behaviors other than optomotor responses were unaffected by stationary patterns (**Figure 1g-n**).

**Figure 4.**
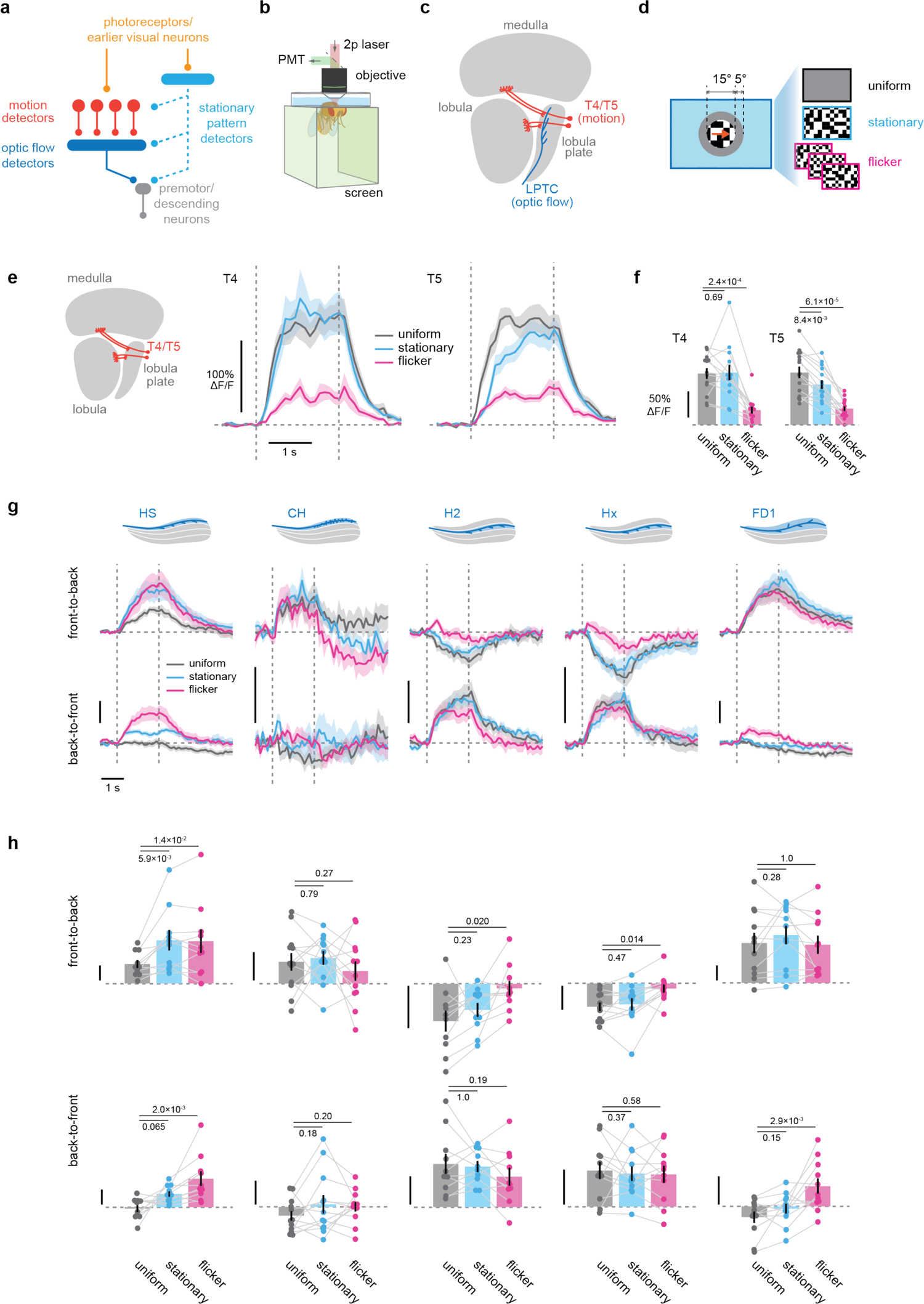
Stationary patterns do not suppress local motion and optic flow detector neurons. **(a)** A schematic of our hypothesis: hypothetical stationary pattern detectors can suppress optomotor responses by inhibiting local motion detectors, optic flow detectors, or premotor neurons. **(b)** A schematic of our two-photon imaging setup. **(c)** In the visual system of *Drosophila* flies, motion is first detected by the elementary motion detector neurons T4/T5. Projection neurons collectively called lobula plate tangential cells (LPTCs) achieve selectivity to specific optic flow patterns by pooling signals from T4 and T5 over space. LPTCs send axons to the central brain, where they likely synapse onto premotor circuitry. **(d)** The modified island of motion stimuli for T4/T5 recording. Background patterns moving at 40 °/s were shown through a circular aperture with 15° diameter, centered around the receptive field of T4 or T5. A 5° wide, gray annulus separated the aperture and the foreground. A slower velocity than the stimuli used for the behavioral experiments were used, because T4/T5 are tuned to slower velocities than optomotor responses^24^. **(e, f)** Calcium responses of T4/T5 neurons to the island of motion stimuli moving in their preferred direction, (e) over time or (f) time averaged, by the different foreground conditions. The vertical dotted lines mark the onset and offset of the stimuli. The horizontal dotted line indicates the pre-stimulus baseline. The responses were averaged over a 3 second window starting at the stimulus onset. See Figure S3a, b for their response to stimuli moving in the non-preferred directions. **(g, h)** Calcium responses of the 5 LPTC types to the islands of motion stimuli moving in the (*top*) front-to-back or (*bottom*) back-to-front directions, either (g) over time or (h) time averaged. Vertical scale bars each indicate 20% ΔF/F. Also see Figure S3c for their response to the foreground patterns without islands of motion. (e, f) T4: N = 13 flies. T5: N = 15 flies. (g, h) HS: N = 10 flies. CH: N = 12 flies. H2: N = 10 flies. Hx: N = 11 flies. FD1: N = 11 flies. Numbers over plots indicate p-values from two-sided Wilcoxon signed-rank tests.

### Stationary patterns do not suppress optic flow detectors

The observation that stationary patterns do not suppress T4 and T5 suggests that information about stationary patterns is integrated into the circuitry for optomotor response somewhere downstream of these neurons. T4 and T5 neurons synapse onto several types of wide-field, optic-flow sensitive projection neurons collectively called lobula plate tangential cells (LPTCs)^43–47^ (**Figure 4c**). LPTCs tuned to horizontal optic flow are thought to implement optomotor responses, although only HS neurons have been causally connected to optomotor behavior thus far^48–50^. Here, we recorded calcium responses of five different types of putatively yaw tuned LPTCs: HS^12, 45^, H2^15, 47, 51, 52^, CH^15, 43, 47, 53^, Hx^54^, and FD1^47, 55^ (**Figure 4g, h**). We imaged the lobula plate dendrites of these cells expressing jGCaMP7b^56^, which has a relatively low Ca^2+^ dissociation constant that is suited for detecting neural hyperpolarization. As visual stimuli, we used the islands of motion stimuli with the uniform, flicker, and stationary configurations used in the behavioral experiments, as well as full-field stationary, flickering, and translating checkerboards. All the LPTC types studied exhibited directionally selective responses to moving checkerboards, with preference to either front-to-back (HS, CH, FD1) or back-to-front (H2, Hx) directions (**Figures 4g, h, S3d**), consist with the lobula plate innervation^43, 47, 54^ and previous physiological recordings^12, 15, 45, 54, 55^. Negative calcium responses to motion in their non-preferred direction were also detected in some LPTC types (CH, H2, Hx) (**Figures 4g, h, S3d**). Overall, stationary foreground patterns did not affect the responses of LPTCs, except for significantly increasing HS responses (**Figures 4h**). Thus, these calcium imaging experiments show that stationary patterns suppressed neither T4/T5 nor various LPTC types, suggesting that the suppression of optomotor turning signals likely occurs deeper in the brain.

### Evidence for and against self motion are integrated across the eyes

Overall, the imaging results favor the idea that integration of evidence for (i.e., optic flow) and against (i.e., stationary patterns) self rotation takes place in the circuitry downstream of the LPTCs, somewhere in the central brain, rather than in the optic lobe. To further test this hypothesis, we next checked whether the suppression of optomotor response by stationary patterns can transfer between the eyes. To do so, we devised a set of stimuli where different visual patterns (i.e., uniform gray, stationary, flickering, or rotating checkerboards) were presented on each eye of the fly (**Figure 5a**). The two patterns were separated by a 30° wide buffer zone of uniform gray to make sure each eye sees only single type of pattern. Here, if the stationary pattern is suppressing signals that drive optomotor response independently in each optic lobe, stationary patterns presented on the contralateral eye would not suppress optomotor response triggered by rotating stimuli presented on the ipsilateral eye (**Figure 5b**). Conversely, if stationary patterns suppress signals driving optomotor response centrally after information from both optic lobes is integrated as we suspect, then stationary patterns presented on one eye should suppress optomotor response caused by visual motion presented on the other eye (**Figure 5b**). We found that, after accounting for the flies’ innate attraction towards spatial patterns over uniform gray (**Figure 5c**), stationary patterns presented on one eye significantly reduced optomotor turning caused by motion on the other eye when motion pointed the back-to-front direction (**Figure 5d**). In contrast, flickering patterns presented on one eye never suppressed optomotor response triggered by motion on the other eye (**Figure 5d**). Rather, flickering patterns on one eye increased turning caused by front-to-back motion on the other eye (**Figure 5d**). These observations suggest that the integration of information about stationary patterns into the optomotor pathway occurs centrally, where information from the two optic lobes is integrated. In contrast, flickering patterns likely suppress the rotational optomotor response by suppressing flow-sensitive visual neurons locally in each optic lobe, as previously measured ^23, 57^.

**Figure 5.**
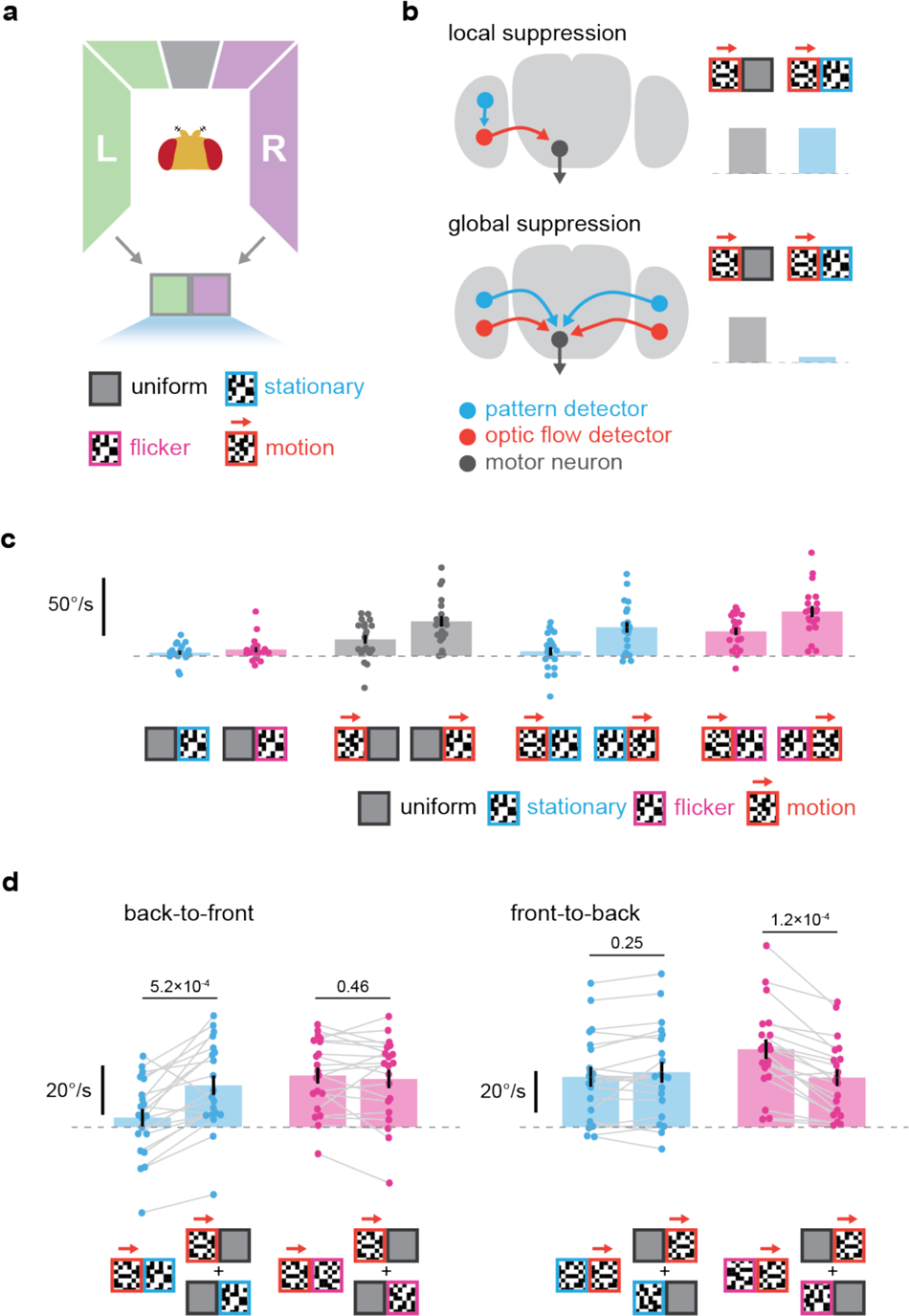
Suppression of rotational optomotor response by stationary patterns translate across eyes. **(a)** A schematic of the binocular stimuli used in (c, d). Different visual patterns were presented on each eye of the fly. Patterns presented on each eye are indicated by icons with two boxes. The patterns were either uniform grey, stationary or flickering checkerboards, or checkerboards moving horizontally in either direction. **(b)** If stationary or flickering patterns suppress optic flow signals in each optic lobe locally, contralaterally presented patterns should not suppress optomotor responses caused by ipsilaterally presented visual motion. Conversely, if suppression of optomotor responses occurs globally, then contralateral patterns should suppress ipsilaterally triggered optomotor responses. **(c)** Time-averaged turning responses to unilateral motion stimuli paired with different foreground patterns. Note that flies exhibited slight turning towards unilaterally presented stationary or flickering patterns. **(d)** Time-averaged turning response to unilateral visual motion paired with contralateral stationary or flickering patterns (*left bars*), compared with expected turning calculated as linear sums of the turning to unilateral visual motion and unilateral stationary or flickering patterns (*right bars*). N = 20 flies. Numbers over bar plots indicate p-values from two-sided Wilcoxon signed-rank tests.

### Behavioral genetic screening for the stationary pattern detectors

The behavioral and imaging experiments have suggested that the integration of the visual negative evidence into the optomotor circuitry happens downstream of LPTCs. Unfortunately, however, optomotor circuitry downstream of LPTCs is currently only poorly mapped. Therefore, instead of trying to track the optomotor pathway deeper into the brain, we instead employed a behavioral genetic screen to directly identify neurons that contribute to hypothetical stationary pattern detectors. To this end, we silenced synaptic outputs of candidate neurons by introducing a temperature sensitive defective allele of *shibire*^58^ using the Gal4/UAS system, and repeated the behavioral experiments with the islands of motion stimuli. We then looked for Gal4 driver lines whose silencing rescued fly optomotor response in the presence of stationary patterns. The degree of suppression was quantified as the ratio between the flies’ time-averaged turning amplitude in response to the islands of motion stimuli with stationary and uniform foregrounds (henceforth fractional turning). This value would be 0 for complete suppression of turning by a stationary foreground and 1 for no suppression at all. For the screen, we selected single and split Gal4 lines targeting the following neuron classes: columnar neurons in lamina, the most peripheral neuropil of the fly visual system (L and C cells)^44, 59–61^, columnar input neurons to the motion-detecting T4/T5 cells (Mi and Tm cells)^44, 62–64^, multi-columnar, amacrine-like interneurons in distal medulla (Dm cells)^44, 65^, and neurons in the heading circuitry and their visual inputs (E-PG, Ring, and TuBu cells)^66–68^. Note that, although these drivers were selected based on their expression in specific visual neuron types, their expression patterns are not strictly limited to these neurons but rather can be broader. After the initial screening, to exclude false positives due to the large number of statistical comparisons, we repeated the identical behavioral experiment a second time on the lines whose silencing resulted in significant changes in behavior.

Among 37 driver lines we screened, we identified four Gal4 lines whose silencing reproducibly resulted in significantly reduced optomotor suppression by stationary patterns. One of the drivers was a split Gal4 line (SS00316) designed to selectively label Mi4 neurons^69, 70^ (**Figures 6a, b, S4c**). The other three drivers were single Gal4 drivers, R13E12, R15C05, and R75H07 (**Figures 6a, b, S4c**). Although these drivers were selected for their expression in visual neurons (R13E12 for Tm3, R15C05 for Dm4, Dm9, Dm12, and R75H07 for Dm4, Dm11, L3)^65^, silencing the same set of visual neurons with independent drivers did not result in the same phenotype (R55D08 for Tm3, R23G11 for Dm4, SS02427 for Dm9, R11C05 for Dm11, R47G08 for Dm12, split L3 for L3) (**Figure 6a**), implying that “off-target” labeling drove the silencing phenotype. Indeed, the single Gal4 drivers in the hits labeled a large number of neurons in the central brain^71^, and it was difficult to identify which specific cell types contributed to the behavioral phenotype. In an attempt to narrow down specific neuron types labeled by these drivers that contributed to the behavioral phenotype, we created three split Gal4 drivers by combining split Gal4 hemidrivers under the control of the R13E12 enhancer segment and three neurotransmitter identity marker genes (ChAT, vGlut, GAD1)^72–74^ (**Figure S4a**). Among the generated split Gal4 driver, only silencing of R13E12 x ChAT resulted in significant reduction in optomotor suppression by stationary patterns (**Figure S4b**). This result suggests that cholinergic neurons among R13E12+ neurons contributed to the detection of stationary patterns or suppression of optomotor response. However, unfortunately, the expression patterns of these split drivers were still too widespread to identify specific cell types. Importantly, silencing neurons in the heading direction circuitry^66–68, 75^ did not result in significant reduction in the optomotor suppression (**Figure 6a**), suggesting that the suppression is independent of that circuit. Thus, our screen left us with the neuron type Mi4 as a solidly identified contributor to the putative stationary stimulus detector.

**Figure 6.**
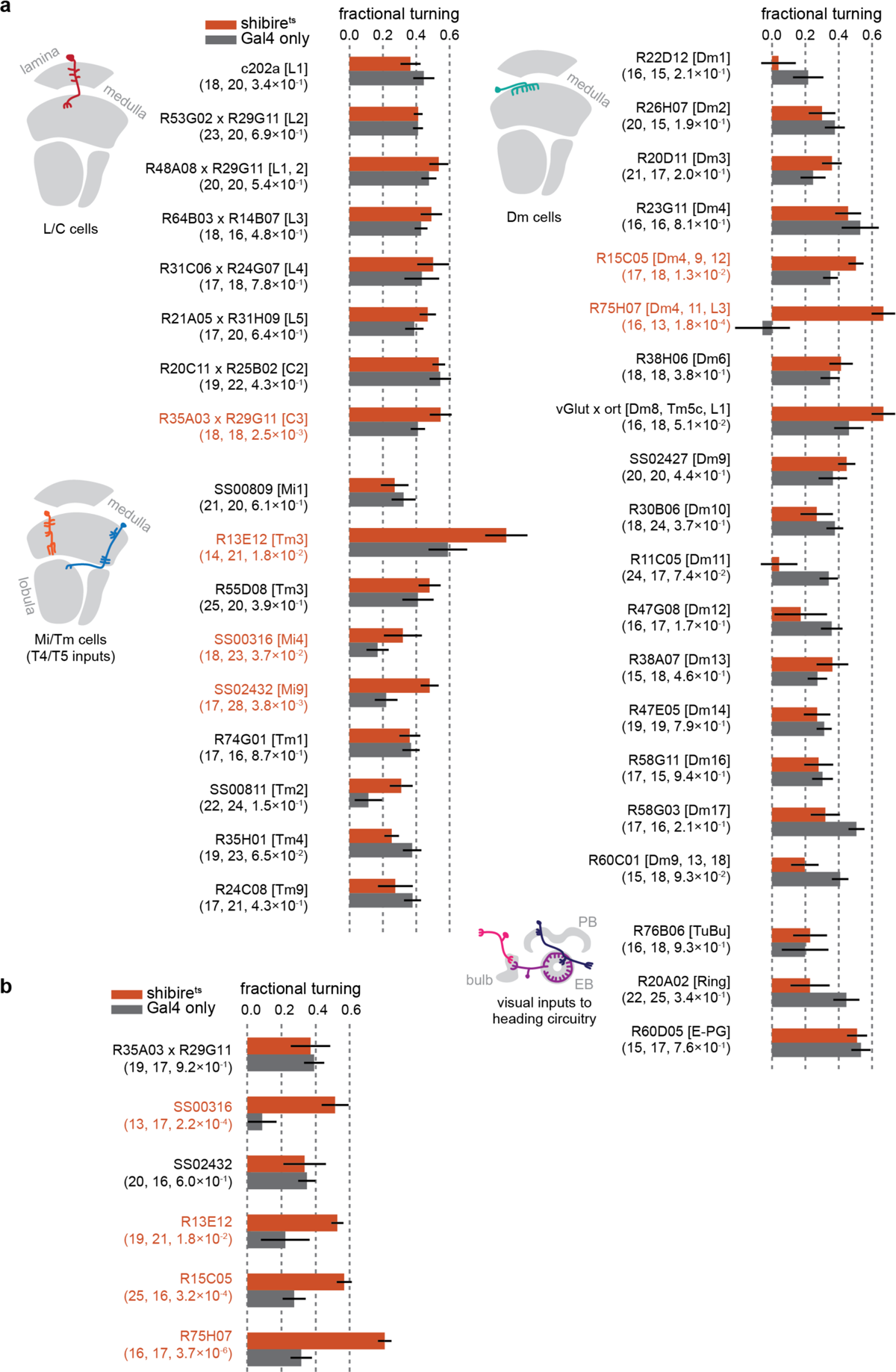
Behavioral genetic screening for stationary pattern detector circuitry. **(a)** The results of the initial screening experiment. The x-axis represents fractional turning, calculated as the ratio between time-averaged turning amplitudes in response to the islands of motion stimuli paired with stationary and uniform foregrounds. Thus, smaller numbers indicate stronger suppression of turning. Orange and gray boxes respectively represent data from flies with both Gal4 and UAS-shibire^ts^ and corresponding Gal4 only controls. The bar plots indicate the mean over the population, and the error bars are standard error of mean. Numbers in the parentheses respectively indicate the sample sizes for flies with and without UAS-shibire^ts^, and p-values from two-sided Wilcocon rank sum tests. The names of the nominal target neuron types are indicated in square brackets. The schematics on the left are showing the approximate neuromorphology of the nominal target neuron classes. Labels for lines that showed statistically significant differences are marked in orange. **(b)** Same as (a), but for the replication experiments. AOTU: anterior optic tubercule; PB: protocerebral bridge; EB: ellipsoid body.

### Response properties of Mi4 can support detecting stationary patterns

Through our screening experiments, Mi4 was identified as necessary for the suppression of optomotor responses by stationary patterns (**Figure 6**). Mi4 is a type of columnar, GABAergic, ON-preferring neuron primarily known for being presynaptic to T4^63, 69, 76^. We therefore considered how responses of Mi4 could support the detection of stationary visual patterns. To this end, we recorded the axonal calcium activity of Mi4 at layer 10 of medulla (M10) using jGCaMP7b, while the flies were presented with full-field checkerboard patterns, either stationary or flickering. On average, Mi4 exhibited only small average responses to both stationary and flickering checkerboard patterns (**Figure 7a**). However, the checkerboard patterns in each Mi4 receptive field could be light or dark in different trials, since checkerboard patterns were chosen randomly. Thus, the response of Mi4 to the stimuli was heterogeneous across trials and ROIs. When we calculated standard deviation of Mi4 responses across trials and ROIs for each time point, we found higher standard deviation for the stationary condition (**Figure 7b**), implying that Mi4 exhibited more varied responses to stationary than to flickering stimuli. To better understand different patterns of Mi4 responses to the stationary checkerboard stimuli, for each fly and stimulus condition, we sorted the trial-by-trial response time traces of Mi4 recordings by their time-averaged response amplitude. We then averaged the sorted time traces within each 20^th^ percentile, and averaged across flies. The analysis revealed that Mi4 exhibits both positive and negative sustained responses to stationary checkerboard patterns (**Figure 7c**), whose magnitude is significantly greater than its responses to flickering stimuli (**Figure 7d**). The sustained, bipolar responses of Mi4 to stationary checkerboards are consistent with its previously documented slow kinetics^69, 76^, its spatially antagonistic receptive field structure^76, 77^, and its lack of rectifying nonlinearity^20, 77^.

**Figure 7.**
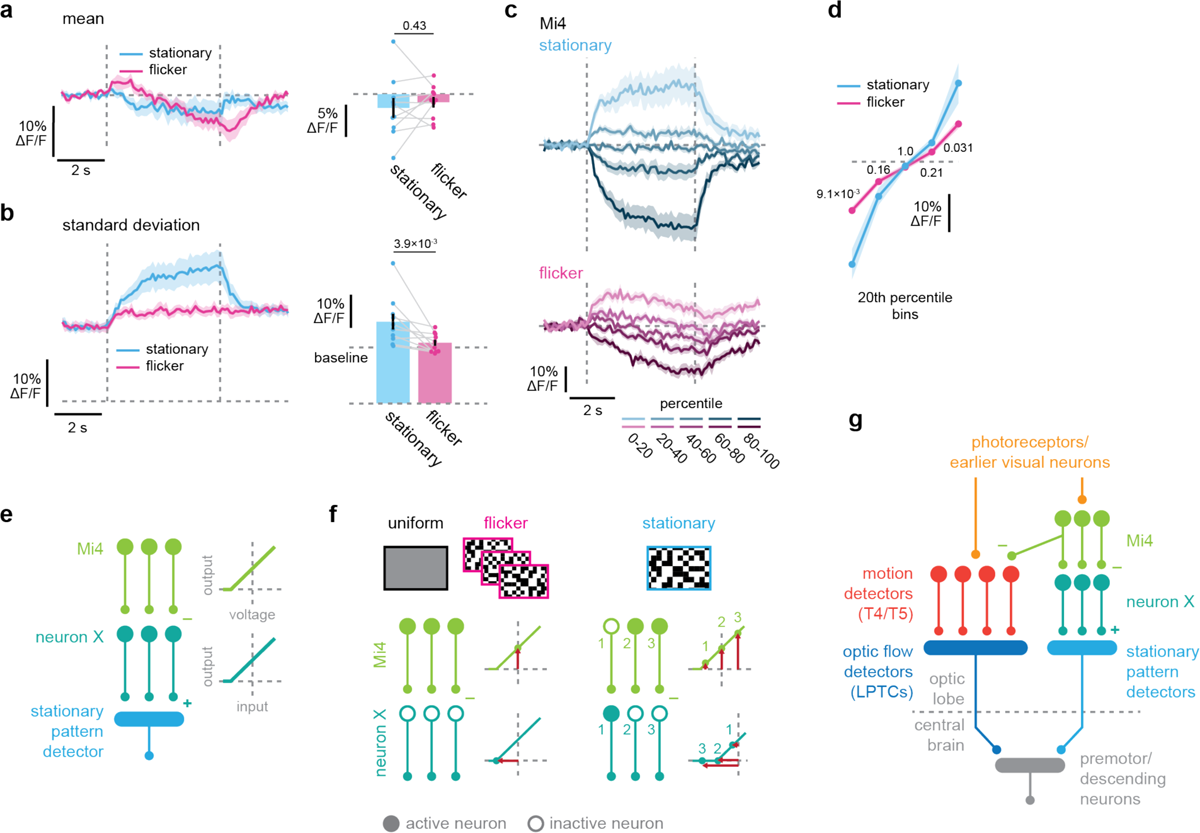
Mi4 is a part of peripheral circuitry that detects stationary patterns. **(a)** Mean calcium responses of Mi4 neurons to the stationary or flickering checkerboards, either (*left*) over time or (*right*) time averaged. The vertical dotted lines mark the onset and offset of the stimuli. The horizontal dotted line indicates the pre-stimulus baseline. **(b)** Instantaneous standard deviation of calcium responses of Mi4 neurons across trial and ROIs by stimulus conditions, either (*left*) over time or (*right*) time averaged. **(c)** The sorted average calcium responses of Mi4 to (*top*) stationary or (*bottom*) flickering binary checkerboards. The trial-by-trial response time traces were sorted by their time-averaged amplitude within each fly disregarding ROI identities, and averaged within each 20^th^ percentile. **(d)** Same as (c), but averaged over time and plotted for each 20^th^ percentile bin by stimulus conditions. **(e)** A schematic of potential downstream circuitry of Mi4 that can detect stationary patterns. (*left*) Here, we assume a layer of columnar “neuron X” downstream of Mi4, whose excitatory outputs are spatially pooled by a downstream stationary pattern detector. (*right*) Input-output relationships of Mi4 and the neuron X. Mi4 is sending non-zero output at the resting potential, indicated by the vertical dotted line. The neuron X also has a positive baseline activity, indicated by the vertical intercept. **(f)** A schematic showing how the neuron X operates. (*left*) When uniform or flickering patterns are shown, activity of Mi4 remains around its resting state, providing inhibitory input to the neuron X. The neuron X remains inactive as a result. (*right*) When stationary patterns are shown, some Mi4 neurons are strongly inhibited by OFF contrast (indicated at position 1), thereby disinhibiting some neuron X (indicated at position 1). Since neuron X outputs are rectified, the spatially averaged activity of the neuron X will be higher when there are stationary patterns. **(g)** A summary schematic of the circuitry involved in the detection of stationary patterns as negative evidence against self rotation and the suppression of optomotor response. Given inputs from earlier visual neurons, T4 and T5 first detect direction of motion within a local receptive field. LPTCs pool the outputs of T4 and T5 over space and send outputs in the central brain. In parallel, visual neurons including Mi4 detect the existence of stationary patterns. Unknown visual projection neurons then send the information about the stationary patterns to the central brain, where commands for optomotor response are suppressed. (a-d) N = 10 flies. Numbers over plots indicate p-values from two-sided Wilcoxon signed-rank tests.

Lastly, we wondered how circuitry downstream of Mi4 could decode the existence of stationary visual patterns. Because of the linearity of the Mi4 receptive field, downstream neurons cannot simply sum Mi4 activity over space to detect stationary visual patterns (**Figure 7a**). Rather, a nonlinear, rectifying operation on the Mi4 outputs seems to be necessary before spatial averaging. For example, imagine a layer of columnar neurons downstream of Mi4 with a positive baseline activity and a non-linear, half-wave rectifying input-output relationship, hereafter noted as the neuron X (**Figure 7e**). Mi4 is a GABAergic, inhibitory neuron, and previous studies have suggested that they have positive baseline activity such that they inhibit their downstream neurons at rest^20, 77^. When uniform or flickering stimuli are presented, Mi4 will remain near its resting state (**Figure 7a**), and their baseline inhibitory output will cancel out with the positive baseline activity of the neuron X, keeping them silent (**Figures 7f**). In contrast, when a stationary visual pattern is presented, some instances of Mi4 neurons will be strongly inhibited by sustained OFF contrast (**Figure 7c**), thus disinhibiting the neuron X (**Figures 7f**). These mechanics tune the neuron X to sustained dark patterns, whose spatially averaged activity can distinguish stationary patterns from uniform and flickering stimuli. A quantitative implementation of this Mi4-based stationary pattern detector model is shown in **Figure S5** (please see also **Methods** > *Stationary Pattern Detector Model* for details). **Figure 7g** summarizes the circuit architecture for the detection of stationary patterns and suppression of optomotor response inferred from the present study.

## Discussion

In the present study, we asked whether and how *Drosophila* exploits negative evidence against self rotation in the face of world motion in order to suppress inappropriate rotational stabilization responses. With psychophysical experiments, we demonstrated that stationary patterns of contrast, especially ones with vertical orientation, selectively suppress flies’ optomotor turning, a heuristic geometrical cue indicating the absence of self rotation (**Figure 1**). In parallel *in silico* experiments, we also showed that ANNs trained to distinguish self and world motion develop a similar behavior as flies (**Figure 2**), supporting the functional interpretation of the fly behavior. Through additional behavioral experiments as well as calcium imaging of neurons in the optomotor pathway, we established that the information about stationary patterns and optic flow are integrated into rotational behavior in the central brain, rather than into low-level motion detection in the optic lobe (**Figures 3-5**). Next, through behavioral genetic screening, we identified multiple genetically defined populations of neurons that are necessary for the detection of stationary patterns and suppression of the rotational optomotor response (**Figure 6**). Finally, we physiologically demonstrated that Mi4, a specific cell type identified in our broad candidate screen, has response properties that allow it to encode stationary patterns (**Figure 7**).

### Functional interpretation of the optomotor response suppression

The rotational optomotor response is generally considered an almost reflexive behavior that counteracts biomechanical instability or environmental perturbations in order to stabilize the heading and gaze of animals. To maintain the course stability, animals should initiate optomotor responses only to the optic flow caused by genuine self motion and not to external, world motion. During genuine observer rotation, visual patterns cannot stay stationary on the retina, and thus the presence of stationary visual patterns can function as negative evidence against self rotation. This is the basic geometrical argument that we use to interpret the suppression of optomotor response by stationary patterns as a strategy to suppress optomotor response in the face of world motion. Additional evidence that supports this functional interpretation are as follows: (1) The suppression was specific to the rotational optomotor response, since stationary patterns did not affect other types of visual motion-dependent behaviors or turning behaviors. (2) The suppression of optomotor response by stationary patterns had an orientation selectivity to vertical patterns, as expected from the geometrical arguments. (3) ANNs trained to distinguish self and world motion also learned to predict low probability of self rotation for stimuli including stationary patterns. Overall, these observations convincingly support the interpretation that the suppression of optomotor response by stationary patterns reflects an evolved strategy to distinguish between self and world motion.

At the same time, we do not suggest that the fly’s strategy to distinguish self and world motion is optimized in the sense that it is the smartest conceivable algorithm. For example, during a genuine self rotation, there is movement not only of boundaries between areas of different luminance but also of boundaries between areas with different motion signals. We observed that stationary borders between areas with and without visual motion did not strongly suppress optomotor response (**Figure S1d-f**), suggesting that flies do not interpret these stationary boundaries as negative evidence to self rotation. There are at least two different, but not mutually exclusive, explanations as to why this might be. First, while stationary contrast patterns are likely widely encountered under natural conditions, it is more difficult to imagine natural cases in which there is a static boundary between moving and stationary regions on the retina. This infrequency could explain why flies have not evolved a strategy to exploit motion-defined stationary contours as negative evidence against self rotation. Second, motion-defined stationary edges might be too complicated to detect within the architectural constraint of the fly optic lobe. Detection of motion-defined stationary contours would require computing spatial derivatives of motion detector outputs, and then integrating them over time. Neuroanatomically, such computation is likely to require at least several layers of retinotopically-resolved visual neurons postsynaptic to T4/T5 in flies. However, the lobula plate, where T4/T5 axons reside, is already one of the deepest retinotopic neuropils, and adding new layers may be evolutionarily or developmentally difficult. The observation that trained ANNs, especially ones with deeper and wider architectures, were able to distinguish stimuli with and without motion-defined stationary edges (**Figure S2d, e**) is consistent with this second explanation.

One caveat of the present study is that all the stimuli were purely rotational. In practice, rotational optomotor responses typically take place while the observer is also moving forward. Thus, what the observer typically experiences is a superposition of translational and rotational optic flows. However, even in this more naturalistic situation, there is a good reason to believe that the stationary pattern-based algorithm to detect the lack of self rotation can function appropriately. This is because distant visual landmarks can effectively remain stationary on the retina of the observer even during forward locomotion. To an observer moving forward at the speed of *v* m/s, a landmark to the side that is *D* m away from the observer will appear to be moving at an angular speed of *v/D* rad/s. For instance, given that typical walking speeds for *Drosophila* are around 2 cm/s^78, 79^ and that the receptive angle of each ommatidia is just below 0.1 radian^19^, an object one meter away would appear to move at the pace of 0.2 ommatidia/s (i.e. 0.2 Hz) to a walking fly. The temporal frequency of 0.2 Hz is an order of magnitude lower than the preferred frequency of change detecting visual neurons in active flies^76^, and thus such object should appear virtually stationary. Thus, stationary visual patterns can likely help walking flies distinguish world motion from genuine self rotation even during forward locomotion.

### Matched filtering and different types of evidence for self motion estimation

Locomotor stability is fundamental to any goal directed behavior, and thus the algorithms animals use to estimate self motion from visual inputs need to be accurate. Existing models of visual self motion estimation typically employ some form of template matching algorithms over vector field representation of visual motion^13, 14, 80, 81^. These algorithms take spatial patterns of local image velocity as inputs, and compare them with optic flow templates during a certain type of self motion (e.g., clockwise yaw). Template matching can thus estimate what type of self motion is most likely given visual motion, assuming that there is indeed self motion. Here, local motion consistent and inconsistent with the flow template are respectively treated as evidence *for* and *against* the specific self motion. In particular, negative weighting of flow-inconsistent local motion is important for distinguishing partially overlapping patterns of optic flow (e.g., forward translation vs. yaw rotation or vertical translation), whose mechanistic implementation has been well studied in insects^15, 16, 46^. However, since these template matching algorithms take local image velocities as their inputs, by design, they treat stationary visual patterns and mere uniformity identically — as zero input — while the two can have quite different implications for assessing self motion. As we discussed above, stationary edges strongly argue against self rotation orthogonal to them, and thus they function as *evidence of absence of self rotation*. In contrast, uniform parts of the visual field are consistent with any true velocity, and thus they are simply *absence of evidence for self rotation*. The fly’s behavior distinguishes between these two different types of zero motion signal, but the canonical template matching algorithms do not. Thus, they must be supplemented to account for our data in the fly. Here, we have provided a simple model framework that explains how a stationarity detector could function and suppress optomotor turning (**Figure 7**).

While we focused on the use of visual cues for estimating self motion, animals have access to additional non-visual cues useful for estimating self motion (or lack thereof). For example, how animals use the sensation of acceleration to stabilize gaze and locomotion has been well studied in both vertebrates and insects^82, 83^. While acceleration sensing is less susceptible to confounding environmental factors such as world motion or illumination, it instead suffers from unique mechanical limitations, like the inability to encode translations with constant velocity and the inability to distinguish translational acceleration from gravity^82^. Another important cue for self motion estimation is the efference copy of motor commands. From the perspective of estimating the net amount of self motion, motor commands can be considered as strong evidence *for* the existence of self motion. In contrast, from the perspective of stabilization behaviors, animals need to estimate *inadvertent* (rather than net) self motion to be corrected. To do so, expected self motion, given the motor commands, should be subtracted from net self motion estimated from sensory inputs^84^. Interestingly, recent studies in *Drosophila* have reported that optic flow-sensitive LPTCs switch between encoding net and inadvertent self rotation depending on their locomotor modes: In walking flies, optic flow and motor commands act additively in both HS^51, 85, 86^ and H2 cells^51^, such that their response amplitudes are maximal when flies are voluntarily turning and receiving congruent visual feedback, consistent with net rotation encoding. In contrast, in flying flies, visual and motor signals act subtractively, such that HS cells are silent during voluntary turns^50, 87, 88^, consistent with encoding inadvertent rotation. It is currently unclear how this switch is achieved, or how downstream circuits decode outputs of LPTCs differently depending on locomotor modes.

A powerful framework to understand how animals may combine different sensorimotor cues to accurately estimate self motion is Bayesian inference^3, 89–91^. In a Bayesian inference framework, the problem of self motion estimation can be formulated as the problem of finding the maximally likely self motion velocity, ν*_self_*, which encompasses both translation and rotation, given its distribution conditional to sensorimotor cues. That is: ν*_self_* = argmax p(ν*_self_*|S), where S represents various sensorimotor cues. Following Bayes’ rule and assuming independence of sensory channels, p(ν*_self_*|S) can be expressed as proportional to the product of modality-wise conditional probabilities such as p(vision |ν*_self_*), p(vestibular |ν*_self_*), p(efference |ν*_self_*), as well as a prior distribution p(ν*_self_*). The logic of falsification, or negative evidence integration, is inherent here, because if a certain cue present is inconsistent with non-zero self motion (*i.e.*, p;cue <ν*_self_* ≠ 0? = 0), that will strongly drive p(ν*_self_* ≠ 0|S) down, bringing ν*_self_* close to 0. Motion-based template matching models in visual self motion estimation describe p(vision |ν*_self_*), which is equated in models to p;optic flow <ν*_self_*? ∝ ∏_x_ p(ν(x)|ν*_self_*), where ν(x) is local retinal motion at retinal position x ^90^. This template matching can treat local motion cues ν(x) that are inconsistent with a certain ν*_self_* as negative evidence, because p;ν(x)<ν*_self_*? = 0 implies p;ν*_self_*<S? = 0 for that ν*_self_*. In Bayesian terms, the central conceptual statement of the present study can be rephrased as that p(vision |ν*_self_*) should not simply be equated to p(optic flow |ν*_self_*). This is because p(stationary pattern|ν*_self_*), which is also a part of p(vision |ν*_self_*), can have a profound impact on the final posterior p(ν*_self_*|S), since p;stationary pattern<ν*_self_*? ≈ 0 when the rotational component of ν*_self_* is non-zero. This is in contrast to lack of visual motion in the optic flow field, which does not imply lack of self motion (i.e., p;ν(x) = 0<ν*_self_*? ≠ 0 for any ν*_self_*), because any ν*_self_* can result in ν(x) = 0 where portions of the scene are featureless.

### Neural circuit for detecting negative evidence against self rotation

Through a series of behavioral and imaging experiments, we constrained the circuitry that detects stationary patterns and suppresses optomotor turning. First, through calcium imaging experiments, we directly demonstrated that elementary motion detectors (T4/T5) and optic flow detecting neurons (LPTCs) are not suppressed by stationary visual patterns (**Figure 4**). The behavioral observations that motion-dependent behaviors other than optomotor response are not suppressed by stationary patterns (**Figure 1g-n**) are consistent with these imaging results. Thus, information about stationary patterns is likely integrated into the circuitry for optomotor response in the central brain, somewhere downstream of LPTCs. The observation that stationary patterns presented on one eye can suppress optomotor response triggered by optic flow presented on the other eye (**Figure 5**) is also consistent with this “central integration” hypothesis. Unfortunately, the circuitry that translates the activity of LPTCs and other lobula neurons into the turning commands remains poorly understood. Recent functional studies have begun to uncover descending neurons that are involved in stimulus-driven turning, such as DNa02^92, 93^. It is of future interest to identify neurons that link LPTCs to these descending neurons, and look for where the signature of suppression by stationary patterns emerges.

Second, through a behavioral genetic screening, we identified several Gal4 drivers whose silencing reproducibly reduces suppression of the optomotor response by stationary patterns (**Figure 6**). The identified drivers included a split Gal4 driver for Mi4, a columnar GABAergic cell type presynaptic to T4^63, 64, 69, 76^. Previous studies have shown that Mi4 has a sustained, ON-center OFF-surround receptive field^69, 76, 77^, which generate sustained responses to stationary visual patterns. Interestingly, the receptive field property of Mi4 resembles the stationary center-surround antagonistic kernels we found in a simple ANN optimized to distinguish self and world motion (**Figure 2e, S2a-c**). However, unlike ANN units, the output of Mi4 is not strongly rectified^77^, and thus one cannot detect stationary patterns based simply on spatially averaged activity of Mi4 (**Figure 7a, S5d**). Rather, an additional layer of retinotopically resolved, rectifying neurons (putatively labeled neuron X in **Figure 7, S5**) would be necessary to detect the presence of stationary patterns based on Mi4 activity (**Figure 7e-g, S5**). In essence, our Mi4-based stationary pattern detector model simply counts the number of neuron X disinhibited by sustained dark spots in the scene (**Figure S5b-g**). This mechanism parsimoniously explains why darker-than-mean stationary patterns more strongly suppress optomotor response (**Figure 3f**), and is similar to how T4 acquires sensitivity to OFF contrasts^20, 77^. Until recently, our knowledge of downstream neurons of Mi4 was limited to T4 and their other inputs, which have axo-axonal contacts with Mi4 (Mi9 and TmY15)^63^. The recently released whole-brain connectome^94^ identified various columnar output neurons of medulla (e.g., Tm16, TmY3, Y3, Tm20) to be downstream of Mi4, which constitute strong candidates for neuron X.

The remaining three hit drivers from the screen (R13E12, R15C05, R75H07) were selected for their labeling of visual neurons (R13E12 for Tm3, R15C05 for Dm4, Dm9, Dm12, and R75H07 for Dm4, Dm11, L3). However, silencing these visual neurons with other Gal4 drivers (R55D08 for Tm3, R23G11 for Dm4, SS02427 for Dm9, R11C05 for Dm11, R47G08 for Dm12, R64B03 x R14B07 for L3) did not result in similar phenotypes (**Figure 6a**). This observation suggests that other neurons targeted by the three hit drivers, most likely in the central brain, contributed to detection of stationary patterns or suppression of optomotor response. Another possibility is that the silencing phenotypes of these hit drivers might be a cumulative effect of silencing multiple cell types. For example, R15C05 and R75H07 both target Dm4^65^, which, incidentally, happens to be upstream of Mi4^94, 95^. Although silencing Dm4 with R23G11 did not result in a similar phenotype, it is conceivable that this was because Dm4 was redundantly necessary for the stationary pattern detection with other R15C05 and R75H07 positive, but R23G11 negative, neurons.

### Geometrical negative evidence against self rotation in different systems

Optic flow-based course stabilization behaviors are fundamental to any goal directed behavior and have therefore been found across diverse taxa. For example, vertebrates ranging from fish^96^ and birds^97^ to humans^98, 99^ use optic flow to stabilize their locomotion, similar to fly optomotor responses. Mollusks, another major phylum with image forming visual systems, also exhibit an analogous optomotor response^100^. In addition to directly affecting locomotion, optic flow also induces conscious perception of self motion to human observers, called visual vection^101, 102^. Since the geometrical rule that visual patterns cannot remain stationary during rotation of the observer is a general constraint of the visual world, we expect that flow-based course stabilization and perception of self rotation might be suppressed by stationary pattern regardless of species. Indeed, previous studies in humans have found that stationary patterns strongly suppress rotational vection^103–105^. Interestingly, these studies have consistently observed that stationary patterns can suppress rotational vection only when they are in the background, behind moving patterns, but not when they are in front of moving backgrounds. While the authors do not offer any functional account for this phenomenon, the sensitivity of vection suppression to depth can be interpreted as an adaptation to the fact that primates have limbs that easily come into the field of view: parts of the body of the observer, like limbs, move with the observer and thus can remain visually stationary even during self rotation, constituting a notable exception from the geometrical argument that stationary patterns imply the absence of self rotation. Thus, animals with visible extremities of the body cannot necessarily use visually stationary proximal objects as negative evidence against self rotation.

Another visually-driven behavior often compared to insect optomotor response is optokinetic nystagmus (OKN)^5^. OKN is a syn-directional eye movement triggered by optic flow, which is found across vertebrates from jawless fish^106^ to humans^107, 108^, as well as in arthropods^109^ and cephalopods^100^. Similar to rotational vection, it has been found that vertical stationary patterns can strongly suppress OKN in humans^110, 111^. However, the functional significance of this OKN suppression is less clear: initiating OKN in response to world motion (i.e., moving objects) will result in fixation on the moving object. It is not difficult to imagine such fixation is an actual function of OKN, rather than being a false positive example, as in the case of whole-body optomotor response.

Overall, by focusing on universal geometrical constraints of optic flow, we found that flies utilize separate stream of visual evidence (i.e., stationary patterns) to gate their optic flow-based stabilization behaviors. This observation exemplifies how the small brain in flies can integrate both positive and negative evidence to act on appropriate scene interpretations in the context of important, innate behaviors. It is of particular future interest to examine how the algorithms and circuit mechanisms of negative evidence detection against self motion generalize to different species, beyond *Drosophila*.

## Acknowledgements

We thank the members of the Clark lab for helpful comments and discussions. RT was supported by the Takenaka Foundation and the Gruber Foundation. NCBM is supported by a CAPES fellowship. DAC and this project were supported by NIH R01EY026555.

## Author Contributions

RT and DAC designed the study. RT and MA acquired data. BAB, BA, and NCBM performed preliminary experiments for the study. RT analyzed data. RT and BZ performed numerical simulations. RT, BZ and DAC wrote the paper.

## Data availability

All data presented in the figures are available from the corresponding author upon request.

## Code availability

Code to analyze the experimental data is available from the corresponding author upon request. The code to train and test the ANN models is available from https://github.com/ClarkLabCode/SelfMotionDetectionML. The code to run the Mi4-based stationary pattern detector simulation can be found here: https://github.com/ClarkLabCode/Mi4Decoder.

## Materials and Methods

### Key Resource Table

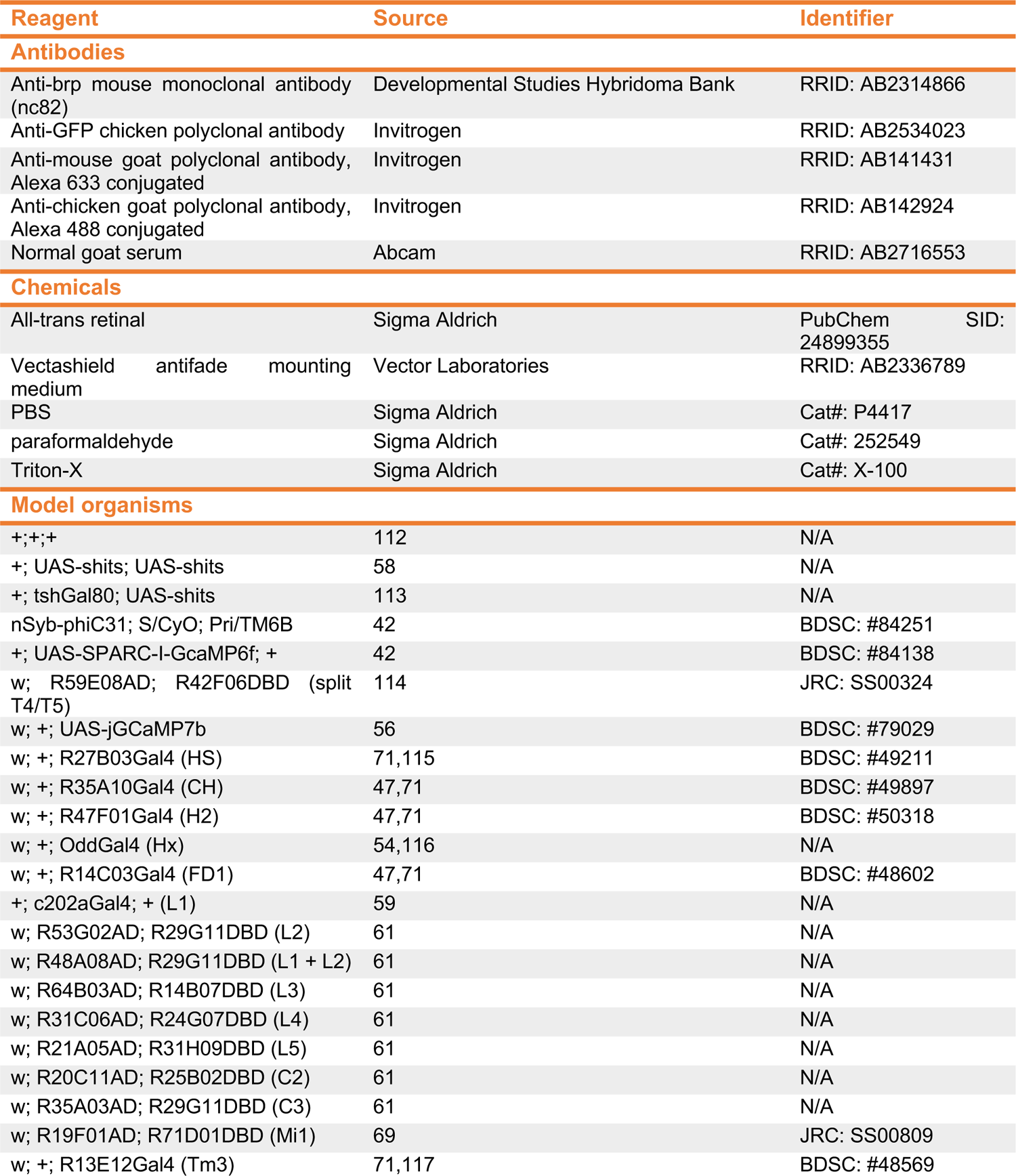
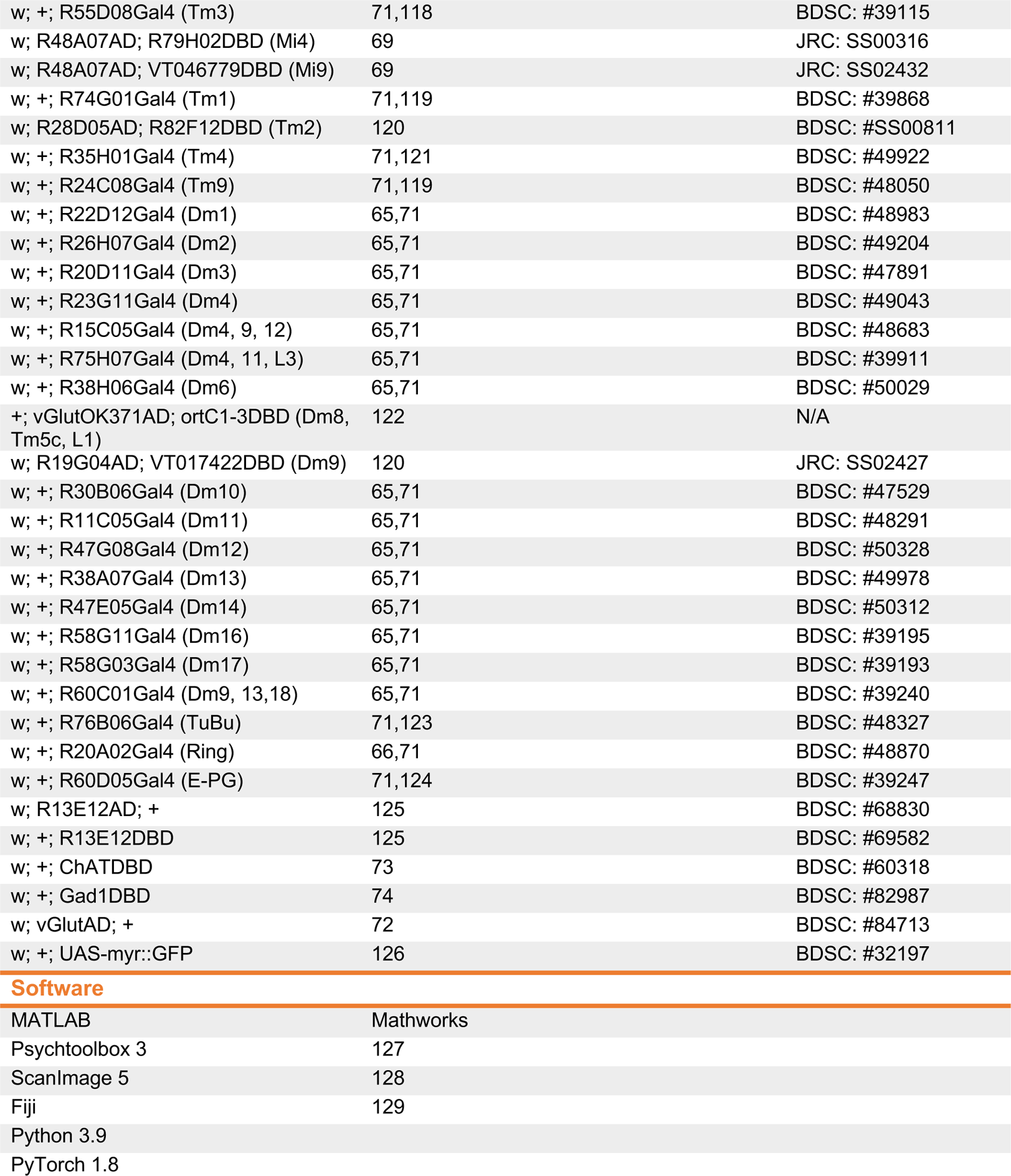

### Fly Strains and Husbandry

All flies were raised at ∼50% relative humidity on a dextrose-based food, under a 12 h:12 h light/dark cycle. Flies for behavioral experiments were raised at 20°C, and flies for imaging experiments were raised at 25°C. Prior to experiments, adult flies were staged on CO_2_ within 24 h post eclosion. All behavioral experiments were performed within 12 to 24 h after staging. For physiology experiments, flies were imaged typically between 2 to 7 days after eclosion. All experiments were performed on non-virgin females. The genotypes of flies used for experiments are compiled in **Table S1**.

### Stimulus Presentation

The stimuli used for behavioral and imaging experiments are respectively compiled in **Table S2, S3**. All stimuli were generated online at 180 Hz. In each experiment, stimuli were presented in a random order, separated by interleave epochs of uniform mean gray, which typically lasted about 3 seconds. In imaging experiments, additional probe stimuli were presented at the beginning and end of the experiment to select responsive regions of interest (ROIs) (**Table S4**). For T4/T5 recordings, we approximately mapped the receptive field location of sparsely labeled T4/T5 axons by interactively presenting translating checkerboards confined in a circular aperture at various locations prior to experiments. Experimental stimuli were then centered about the estimated receptive field center.

### Behavioral Assay and Data Analysis

To measure flies’ visuomotor behaviors, we used a previously reported tethered walking assay (Creamer et al., 2019). Cold-anesthetized flies were tethered onto 30G surgical needles with UV-curable epoxy glue, and mounted on air-floated balls. The flies’ locomotor responses were read out as the rotation of the balls, which we measured with optical mouse chips (resolution ∼0.5°, sampling frequency 60Hz). Visual stimuli were projected onto panoramic screens surrounding the flies using DLP projectors (Texas Instruments Lightcrafter DLP evaluation modules). The visual stimuli covered 270° of azimuth and 106° of elevation. The stimuli were presented only using the green channel of the projectors (peak 520 nm, ∼ 100 cd/m^2^). The behavioral rigs were heated to 36 °C to promote forward walking as well as to use thermogenetic tools (in this study, shibire^ts^).

Flies’ forward walking responses to each presentation of stimuli was normalized relative to the average walking speed within a half second window immediately preceding the stimulus onset. The instantaneous turning velocity as well as normalized forward walking speed were then averaged over trials for each stimulus type to generate individual fly mean traces. Turning and forward walking responses to mirror symmetric pairs of stimuli were also respectively averaged in subtractive and additive fashion. The individual mean traces were then averaged over flies to visualize the dynamics of responses, as well as over time for statistical comparisons across stimulus conditions and genotypes. For the screening experiments in **Figures 6, S4**, flies that exhibited less than 5 °/s turning to the uniform islands of motion stimulus were excluded from the analysis, because small turning amplitude made estimating the degree of suppression of behavior difficult.

### Two-photon Imaging

For imaging experiments, cold-anesthetized flies were mounted onto a holder with UV-curable epoxy. The cuticle on the back of the head, as well as trachea and fat tissue were surgically removed to expose the brain. The brain was then submerged in oxygenated sugar-saline solution^130^. Imaging was performed with a two-photon microscope (HyperScope; Scientifica) with a 20x water immersion objective (XLUMPlanFL; Olympus). Visual stimuli were presented on a panoramic screen surrounding the fly using the same DLP projector as the ones used in the behavioral experiment. The projector light was filtered with a 565/24 in series with a 560/25 filter (Semrock). The stimuli were pitched 45° forward relative to the screen to account for the tilt of the fly head. The stimuli spanned 270° of azimuth and 69° of elevation. The two-photon excitation at 930 nm was provided by a femtosecond Ti-Sapphire laser (Mai Tail; SpectraPhysics). The laser power at the sample was kept below 40 mW. To capture the emission light from green fluorophores and exclude light from the screens, light coming into the photodetector was filtered with two 515/25 filters (Semrock) in series. The microscope was controlled through the ScanImage 5 software^128^ and images were acquired at 8.46 Hz.

### Imaging Data Analysis

ROIs were defined by applying a watershed algorithm^131^ on time-averaged fluorescent images. To remove a small amount of stimulus bleedthrough from the recordings, mean signals from the background region were subtracted from the entire recording. The background region was defined as the largest contiguous region below 10 percentile brightness in the time-averaged image. The fluorescence time traces in each ROI were then transformed into the unit of ΔF/F as follows: The fluorescence within each ROI was averaged across pixels for each frame. The pixel-averaged fluorescence was then averaged over time within each interleave epoch. Decaying exponentials in the form of F = Ae^-t/τ^ were then fit to this averaged fluorescence, where τ was constrained to be identical across ROIs. The assumption is that bleaching of fluorophores should happen uniformly across ROIs. In each ROI, F was subtracted from the original pixel-averaged fluorescence time trace, and reminder (ΔF) was divided by F to generate ΔF/F time traces.

Next, responsive ROIs were selected based on the consistency of their responses to multiple repetitions of probe stimuli presented before the experiments. The response consistency was measured as the average Pearson correlation between all pairs of probe responses. The threshold for response consistency was set at 0.4. For T4/T5 recordings, additional ROI selection criteria were devised to functionally identify specific subtypes of T4 and T5 whose receptive field was appropriately aligned with the visual stimuli. First, in T4/T5 recordings, probe stimuli were repeated at the end of the experiment as well. We discarded T4/T5 ROIs with correlation between the trial-averaged pre-experiment probe response and post-experiment probe response lower than 0.4 as being unreliable. Second, to find ROIs aligned with the stimulus location, we selected ROIs whose direction- and time-averaged responses to the island of motion probe was more than two times larger than their responses to the sea of motion probe (see **Table S4** for definitions of the probe stimuli). Third, to make sure that ROIs belonged to the subtypes of T4/T5 tuned to horizontal motion, we discarded ROIs that responded the most to either the upward or downward island of the motion probe among the four directions. Finally, we selected ROIs based on their direction and polarity selectivity, calculated based on their responses to the short edge probe (see **Table S4**). Direction and polarity selectivity indices (DSI and PSI) were defined as DSI = (r_ftb_ – r_btf_) / (r_ftb_ + r_btf_), PSI = (r_ON_ – r_OFF_) / (r_ON_ + r_OFF_), where r_ftb_ = r_ftb,ON_ + r_ftb,OFF_, r_btf_ = r_btf,ON_ + r_btf,OFF_, r_ON_ = r_ftb,ON_ + r_btf,ON_, r_OFF_ = r_ftb,OFF_ + r_btf,OFF_. R_d,p_ was calculated as the difference between 99 and 50 percentiles of trial-average response time traces to the short edge with direction d (front-to-back (ftb) or back-to-front (btf)) and polarity p (ON or OFF). We labeled ROIs with DSI > 0.4 and PSI > 0.4 as T4a, ones with DSI < −0.4 and PSI > 0.4 as T4b, DSI > 0.4 and PSI < −0.4 as T5a, and DSI < −0.4 and PSI < −0.4 as T5b.

For each stimulus presentation, the average baseline ΔF/F within a half second window immediately preceding the stimulus onset was subtracted from the response time traces. Responses were then averaged over trials for each stimulus type, then over ROIs, to generate individual mean time traces. The individual mean time traces were then averaged either across flies to visualize the response dynamics, or over time to make statistical comparisons between stimulus conditions. Responses of T4 and T5 subtypes were averaged within each fly after appropriately flipping the stimulus directionality. For the Mi4 imaging experiment (**Figure 7c, d**), for each fly and each stimulus condition, by-trial, by-ROI response time traces were sorted according to their time-averaged amplitude ignoring the ROI identities, and averaged within each 20^th^ quantile bin, and then averaged across flies.

### Immunohistochemistry

The brain was extracted from the head capsule in PBS, and then fixated for approximately 15 minutes in 4% paraformaldehyde at room temperature. After three washes in PBS for at least 20 minutes, the brains were blocked with 5% normal goat serum for another 20 minutes. The brains were then incubated with primary antibodies (mouse anti-Brp, 1:25; chicken anti-GFP, 1:50). After another 3 washes, the brains were incubated with secondary antibodies (anti-mouse AF633 and anti-chicken AF488, 1:300). The antibodies were dissolved in PBS with 0.2% Triton-X and 5% normal goat serum. The incubation periods were between 1 to 3 days. The brains were then mounted on glass microscope slides with the Vectashield mounting medium. The Z-stacks were acquired with Zeiss LSM880 confocal microscopes with 20x air or 40x oil-immersion objectives, with a typical slice thickness of 1 μm.

### Artificial neural networks and training

We built a series of artificial neural networks (ANNs) with a convolutional architecture and trained them on natural scene movies to infer whether a movie was produced due to self motion or world motion. PyTorch 1.8 running on Python 3.9 was used for the simulation. The input layer of the ANNs was 30 x 1 x 72, with the first dimension indicating depth (time), the second dimension height, and the third dimension width. Following the input layer, the model had *L* 1 x 72 convolutional layers with each layer having *C* channels (**Figure 2a**). The sizes of the convolutional kernels were 30 x 1 x 3 between the input and the first convolutional layer and *C* x 1 x 3 elsewhere. For each convolutional layer, the input was padded periodically and the rectified linear unit was used as the activation function. The output of the all units in the last convolutional layer were averaged with spatially uniform weights and logistic-transformed to calculate the probability that the given input movies were generated by self motion rather than world motion. Different channels in the last convolutional layer were weighted differently in the output layer. *L* and *C* were varied between 1 to 6 and 1 to 8, respectively. The loss function was the standard cross entropy loss for binary classifications, and the training was repeated 100 times for each *L, C* combination with random initialization. The performance of the trained ANNs were tested with held-out datasets.

To generate the natural scene movies, 241 natural scene images were obtained from an online database ^36^. Each image had a dimension of 251 x 927 and captured a panoramic natural environment, covering 97.5 degrees vertically and 360 degrees horizontally. For the self motion movies, each image was convolved with a 2-dimensional Gaussian kernel with a full width at half maximum of 5 degrees, mimicking the acceptance angle of one ommatidium of a fly eye. To generate the self motion movies, an image was shifted horizontally according to a positional trace, which has been generated by integrating a velocity trace. The velocity trace was simulated as an Ornstein–Uhlenbeck process with an autocorrelation time of 0.2/ln2 s and a standard deviation of 100 °/s, approximately matching typical turning velocities and timescales of flies^38, 79^. The positional trace was 300 ms long in time and had a temporal resolution of 10 ms, and thus, each movie had 30 frames. After this, each frame of the movies was subsampled to be 20 x 72 such that each pixel covered roughly 5 degrees in the angular space. In this way, each movie was a 30 x 20 x 72 volume. The input to the ANNs was a random horizontal slice from this volume, with the size of 30 x 1 x 72. For the world motion movies, we first randomly selected a horizontal slice with a height of 5 degrees from a stationary natural scene image. Next, up to 10 patches with the height of 5 degrees were randomly selected from the same image to serve as the objects and placed randomly on the horizontal slice. The width of each object was randomly sampled from the interval of 5 degrees to 36 degrees. After this, the whole image was convolved with a 2-dimensional Gaussian kernel just as in the self motion cases. The motions of all the objects in each movie are coherent, and the velocity traces were generated the same way as in the self motion movies. The world motion movies were subsampled also in the same way as in the self motion ones, and the horizontal slice that contained the objects were selected as the input. Each natural scene movie was rescaled to have zero mean and unit variance. We generated 144,000 samples for training and 48,800 samples for testing.

To compare the behavior of the training models with those of the flies, we generated the islands of motion stimuli similar to what was presented to flies, as well as full-field binary checkerboards translating horizontally (**Figure 1c, 2f**). The spatiotemporal parameters of the stimuli were kept identical (resolution of 5°, window size of 15°, window coverage of 20%, velocity of 80 °/s), except that the stimuli here lacked the vertical spatial dimension and had the temporal sampling rate of 100 Hz to match the size of the ANN input layer. For each category of the stimuli (full-field, uniform, stationary, or flicker) 2000 different movies were generated, and fed into each of the trained models. We then computed the averages of predicted probabilities and their logit transformations. For each model, 25^th^ percentile, median, and 75^th^ percentile performances over 100 initializations are visualized in **Figure 2g, h, S2e, f**.

### Stationary pattern detector model

We performed a quantitative simulation of a mechanistic model of a hypothetical stationary pattern detector based on Mi4 as follows. First, visual stimuli were generated at the spatiotemporal resolution of 1° and 100 Hz. Stimuli were either uniform grey, a stationary checkerboard (5° checker size), a flickering checkerboard (5° checker size, updated at 15Hz), or a checkerboard rotating in yaw direction (5° checker size, 80 °/s). Thirty instances of each stimulus category were generated. To simulate the ommatidial resolution of *Drosophila*, the stimuli were then convolved with a spatial Gaussian filter with full-width at half-maximum of 5.7°, and downsampled to 5° resolution^19^. Next, to simulate responses of Mi4, a temporal and spatial filter were convolved with the downsampled stimuli. The temporal filter had a shape of *k(t)* = ate^-t/^τ, where τ = 300 *ms*. The spatial filter had the difference-of-Gaussian shape described as 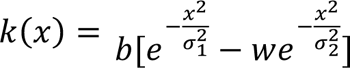, where σ_*_ = 4°, σ_+_ = 8°, *w* = 0.8. *a* and *b* are constants determined so that k(t) and k(x) had unit L1 and L2 norms, respectively. The output of neuron X, a hypothetical downstream neuron of Mi4, is modeled as 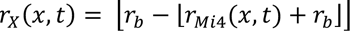, where r*_Mi4_* is Mi4 activity, r_b_is a baseline activity arbitrarily set to 4, and 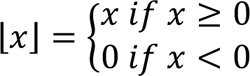. Here, the term 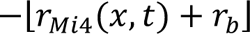 corresponds to GABAergic output of Mi4, which has positive baseline activity. Outer rectification represents synaptic nonlinearity of neuron X. Finally, the response of stationary pattern detector was modeled as the average of r_X_(x, t) over space.

### Quantification and Statistical Analysis

For the statistical purpose, each fly was treated as an independent sample. *P*-values presented are either from double-sided Wilcoxon signed-rank (within-fly, across stimulus condition comparisons) or rank sum tests (across genotype comparisons).

---

**Figure S1.**
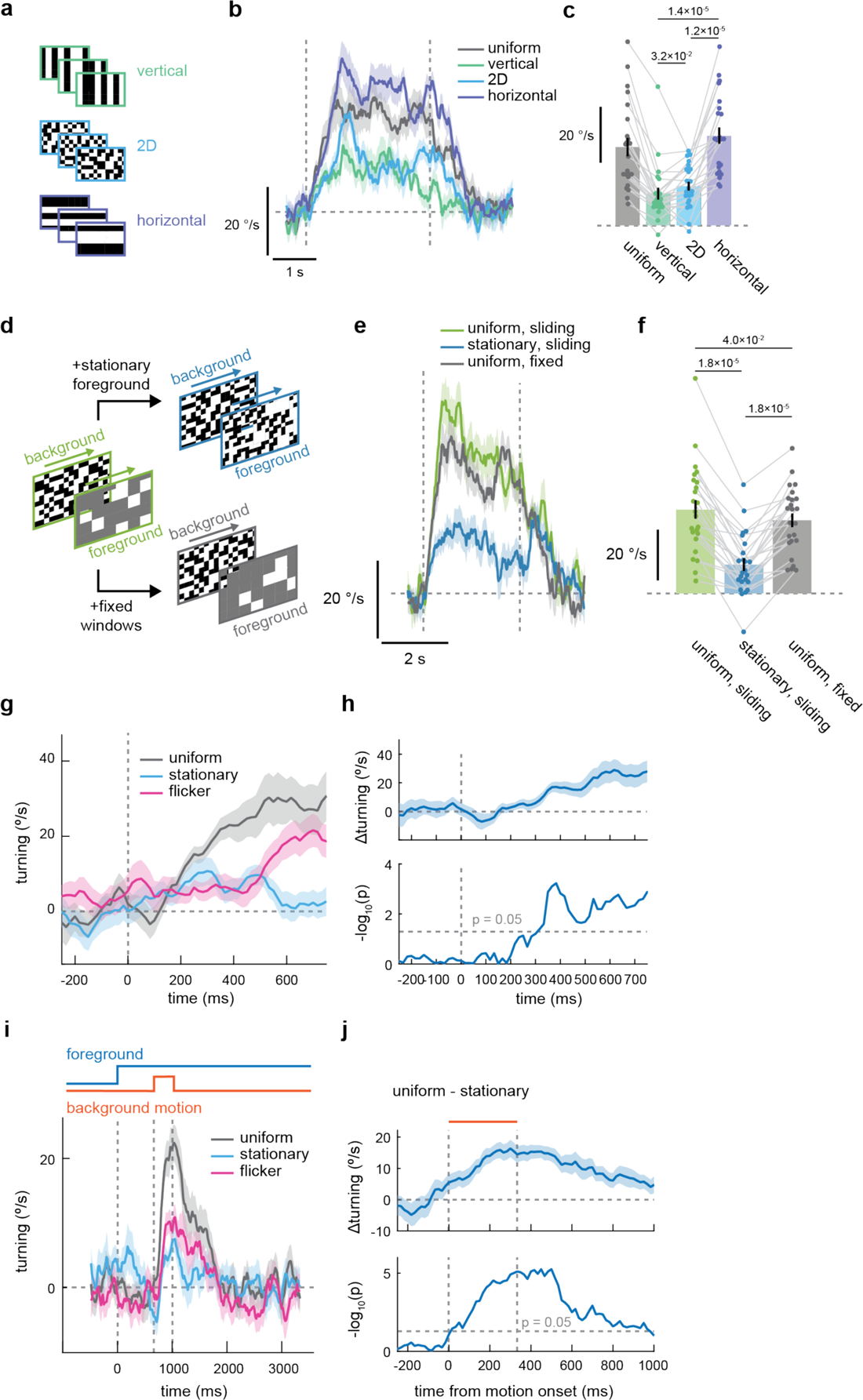
Use of additional geometrical cues to suppress optomotor response. Related to **Figure 1**. **(a)** Schematic of stimuli used to probe orientation selectivity of the optomotor suppression caused by flickering foregrounds. **(b, c)** Fly turning responses to islands of motion stimuli with flickering foreground patterns, either (b) over time or (c) averaged over time, from the same set of flies as in **Figure 1p, q**. **(d)** The islands of motion stimuli with a uniform foreground and windows sliding in the direction and speed of the rotating background (green) is entirely consistent with genuine yaw optic flow. The other two stimuli were created by adding either first-order (contrast-defined stationary patterns in the foreground, not moving with the islands; blue) or second-order (motion-defined stationary contours of stationary windows; grey) cues against self rotation. The stimulus with a uniform foreground and fixed windows is identical to the “uniform” condition in **Figure 1c**. **(e, f)** Fly turning responses to islands of motion stimuli shown in (d), either (e) over time or (f) averaged over time. (g) Same as Figure 1e, but zoomed around the stimulus onset. **(h)** Same as **Figure 1t**, but for the data in **Figure 1e**. (*top*) Difference of turning responses to islands of motion stimuli uniform and stationary foreground patterns. (*bottom*) Negative, log10 transformed p-value between the uniform and stationary conditions, computed at every time point. The horizontal dotted line marks p = 0.05. The p-values reach 0.05 at ∼ 300 ms after the stimulus onset. **(i, j)** Same as **Figure 1s, t**, but with a stimulus where the delay of background motion onset was 2/3 s, and background motion lasted 1/3 s. The p-values reach 0.05 less than 100 ms after the motion onset. (b, c) N = 23 flies. (e, f) N = 24 flies. (g, h) N = 23 flies. (i, j) N = 27 flies. Numbers over bar plots indicate p-values from two-sided Wilcoxon signed-rank tests.

**Figure S2.**
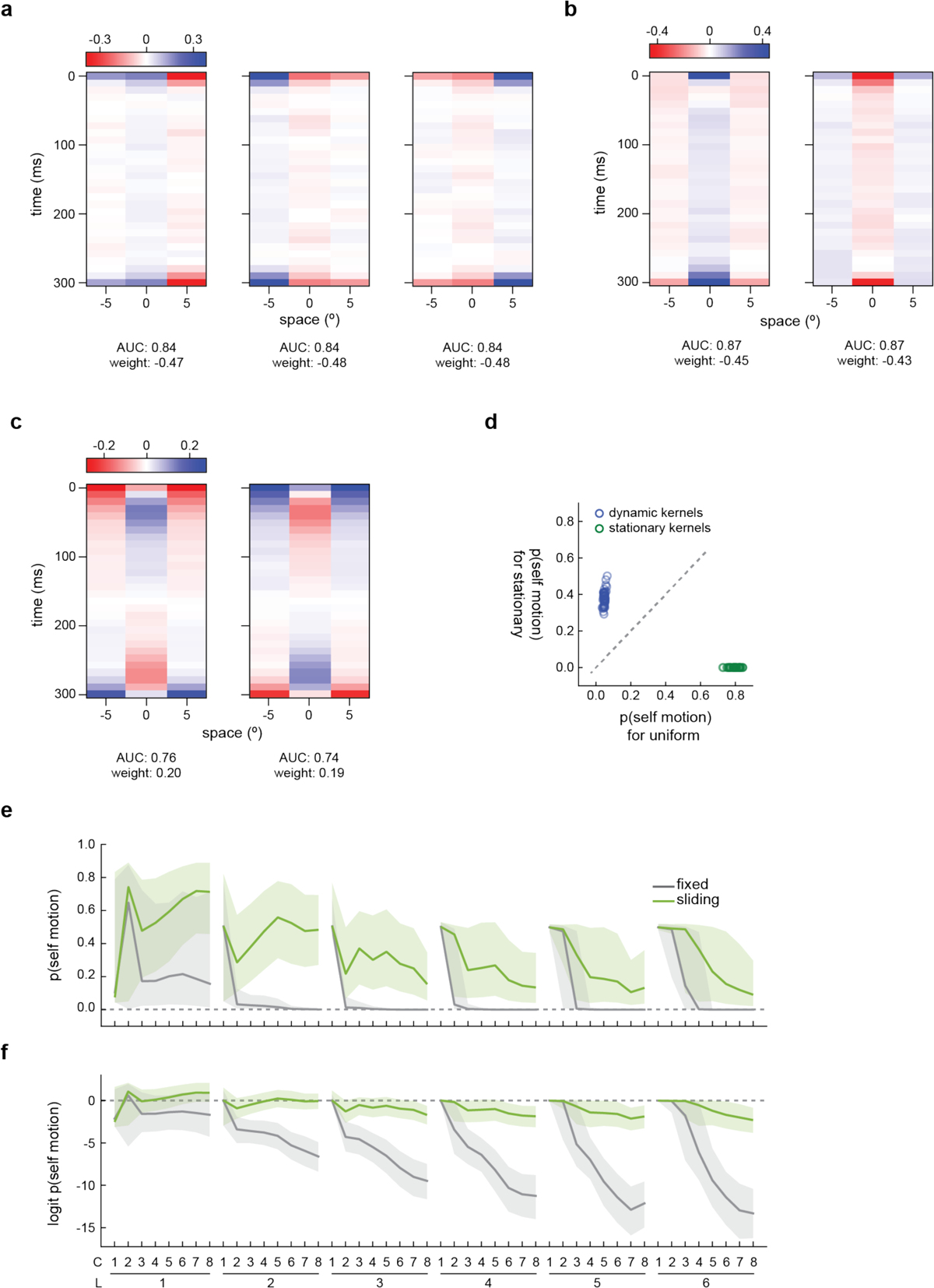
Additional details of the neural network models trained to distinguish self and world motion. (Related to **Figure 2**) **(a-c)** Examples of kernels from single-layer, single-channel models (*L* = 1, *C* = 1), belonging to the category of either (a) stationary edge detectors, (b) stationary center-surround antagonism, or (c) dynamic kernels. Similar to the kernels shown in **Figure 2e**, stationary and dynamic kernels had negative and positive weights toward the probability of self motion, and stationary ones outperformed the dynamic ones. **(d)** The probability of self motion for the islands of motion stimuli with uniform or stationary foregrounds, predicted by each of the 100 trained single-layer, single-channel (*L* = 1, *C* = 1) models. The types of kernels (static or dynamic) are color coded. Models with static kernels, but not ones with dynamic kernels, returned very low probability of self motion for stimuli with stationary foregrounds, analogous to the behavior observed in flies (**Figure 1**). **(e)** The predicted probability of self motion for the islands of motion stimuli with fixed or sliding islands and uniform foreground as shown in **Figure S1d**, as functions of model architectures. The lines represent median, and shaded regions represent 25^th^ and 75^th^ percentile performances across 100 model initializations. **(f)** Same as (d), but showing the logit values instead of the probability of self motion.

**Figure S3.**
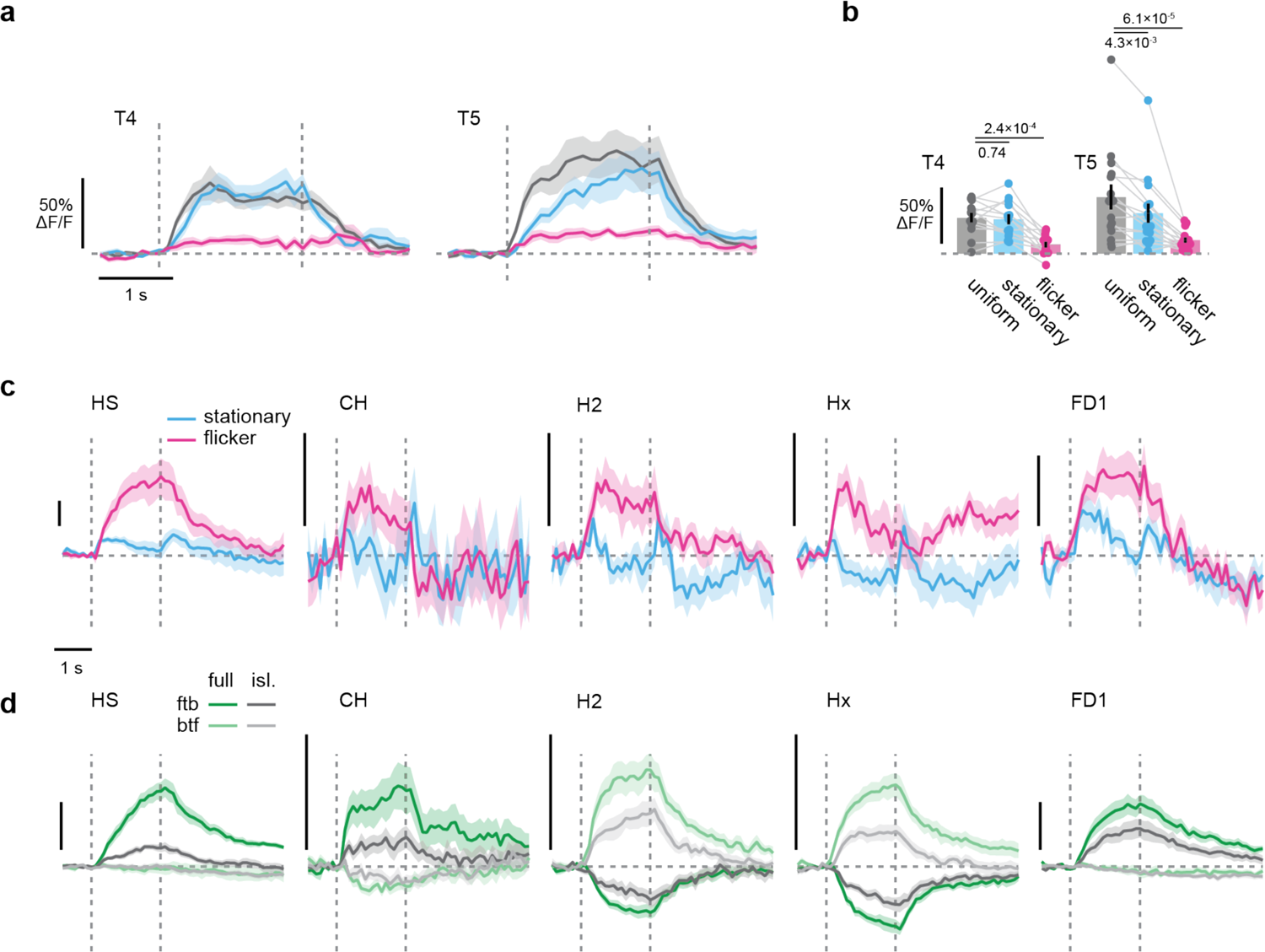
Additional characterization of T4/T5 and LPTCs. Related to **Figure 4**. **(a, b)** Calcium responses of T4 and T5 neurons to the island of motion stimuli moving in their non-preferred direction, (a) over time or (b) time averaged, by the foreground types. **(c)** Calcium responses of the 5 LPTC types to the full-field stationary or flickering checkerboard patterns. Vertical scales bars indicate 10% ΔF/F. Stationary patterns appeared to hyperpolarize H2 and Hx, but these effects were overwhelmed in the presence of islands of motion. **(d)** Calcium responses of the 5 LPTC types to the full-field translating checkerboards (labeled as full) or the islands of motion stimuli with the uniform foreground (labeled as isl.), either moving in the front-to-back (ftb) or back-to-front (btf) directions. Vertical scales bars indicate 50% ΔF/F. (a, b) T4: N = 13 flies. T5: N = 15 flies. (c, d) HS: N = 10 flies. CH: N = 12 flies. H2: N = 10 flies. Hx: N = 11 flies. FD1: N = 11 flies. Numbers over plots indicate p-values from two-sided Wilcoxon signed-rank tests.

**Figure S4.**
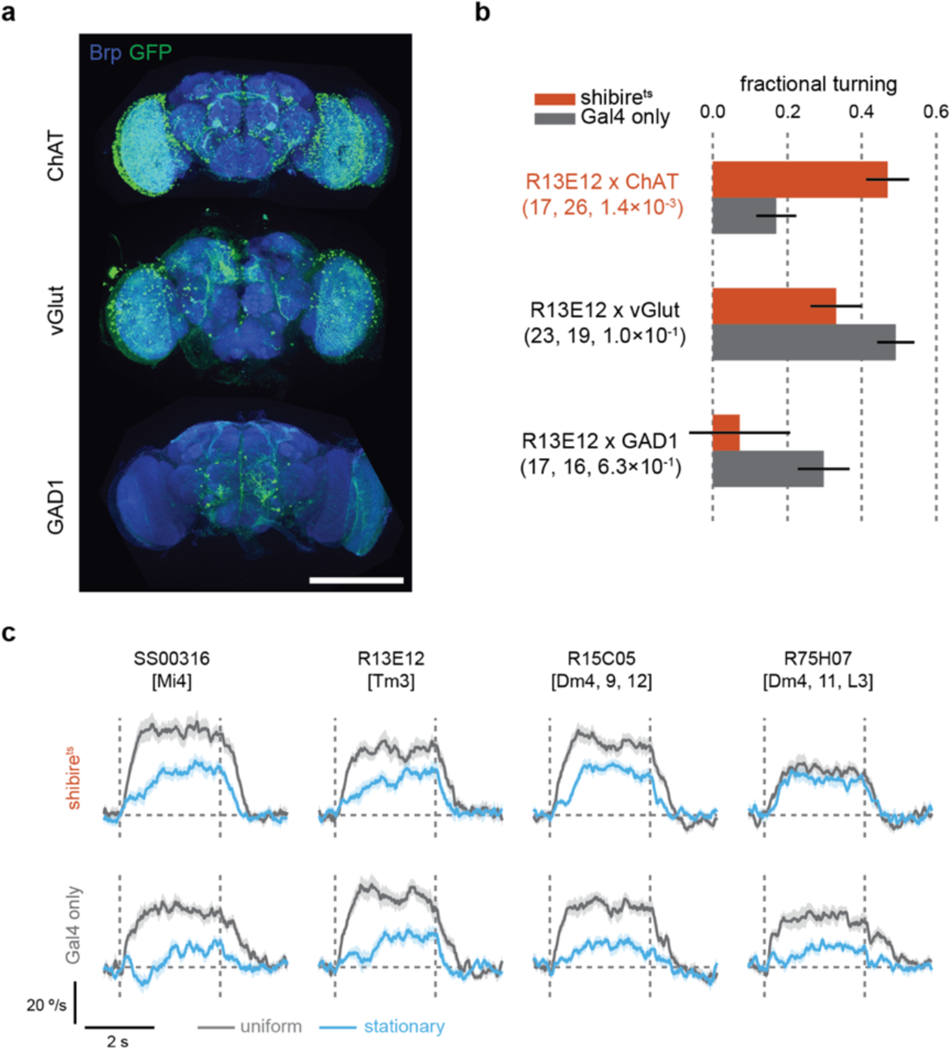
Additional characterization of screening results. Related to **Figure 5**. **(a)** Expression patterns of the split Gal4 drivers between R13E12 and neurotransmitter marker genes. (*top*) R13E12AD x ChATDBD > UAS-GFP, (*middle*) vGlutAD x R13E12DBD > UAS-GFP and (*bottom*) R13E12AD x GAD1DBD > UAS-GFP. The blue and green channels respectively show anti-Brp and anti-GFP staining. The three lines appear to label different sets of neurons. The scale bar indicates 100 μm. **(b)** Fractional turning responses of flies expressing shibire^ts^ under the control of the three split Gal4 lines, similar to Figure 6a, b. Numbers in the parentheses respectively indicate the sample sizes for flies with and without UAS-shibire^ts^, and p-values from two-sided Wilcoxon rank sum tests. Only silencing using the R13E12AD x ChATDBD split driver resulted in a significant rescue of turning in the presence of stationary patterns. **(c)** Raw turning responses for the hit drivers from the replication experiment (**Figure 6b**). Vertical dotted lines mark the beginning and end of the stimuli, and the horizontal dotted line marks zero turning. The difference between the responses to the two conditions is reduced in flies expressing (*top*) shibire^ts^ compared to (*bottom*) the Gal4 only controls.

**Figure S5.**
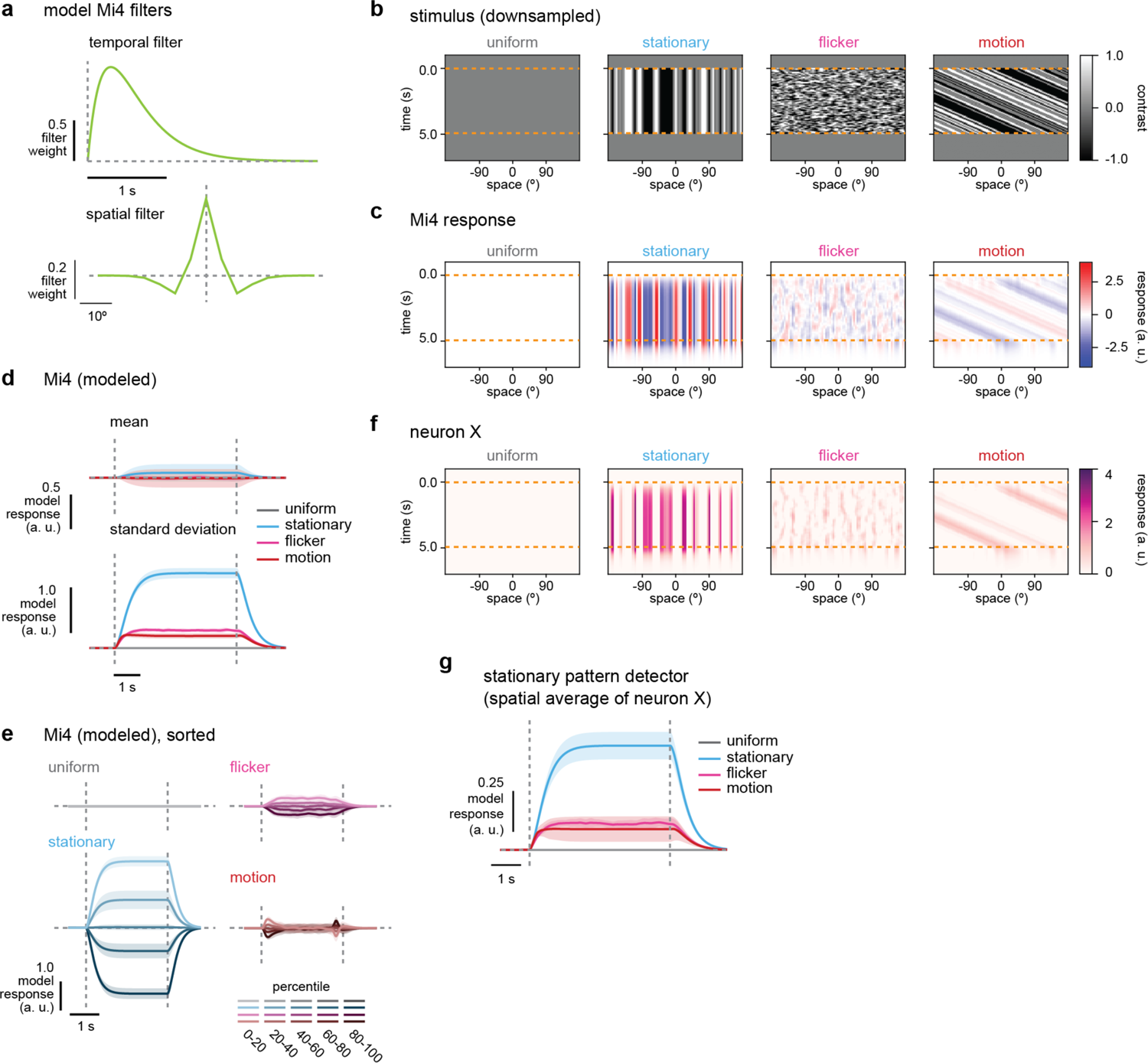
Quantitative simulation of the stationary pattern detector model. Related to **Figure 7**. **(a)** Temporal and spatial kernels of model Mi4, based on previously published measurements^76^. The receptive field of Mi4 is ON-centered, temporally low-pass, and spatially high-pass. See Methods for the details. **(b)** Space-time plots of stimuli presented to the model: uniform grey (“uniform”), stationary checkerboards (“stationary”), flickering checkerboards (“flicker”), and full-field rotating checkerboards (“motion”). Stimuli were generated at the spatiotemporal resolution of 1° and 100Hz, and downsampled to the ommatidial resolution of 5°. Thirty instances of stimuli in each category were generated, and representative examples are shown here. **(c)** The responses of Mi4 neurons, modeled by convolving kernels in (a) with downsampled stimuli in (b). **(d)** The mean and standard deviation of model Mi4 responses across space, as in **Figure 7a, b**. Because of the lack of nonlinearity, on average model Mi4 does not respond strongly to any of the stimuli, while stationary patterns induce larger variability in model Mi4 activity. Shaded area indicates standard deviation across 30 instances of stimuli. **(e)** Sorted responses of model Mi4, averaged across space, as in **Figure 7c**. Similar to the experimental observation, individual model Mi4 strongly responded to stationary checkerboards either positively or negatively, whereas it did not respond strongly to uniform or dynamic stimuli. **(f)** Space-time plots of “neuron X”, as depicted in **Figure 7e, f**. Notice that neuron X is only active when downstream of Mi4 neurons that were inhibited by sustained negative contrast. See Methods for details. **(g)** The response of the hypothetical stationary pattern detector, computed as a spatial average of neuron X activity. Shaded area represents standard deviation across 30 instances of stimuli.

**Table S1.**
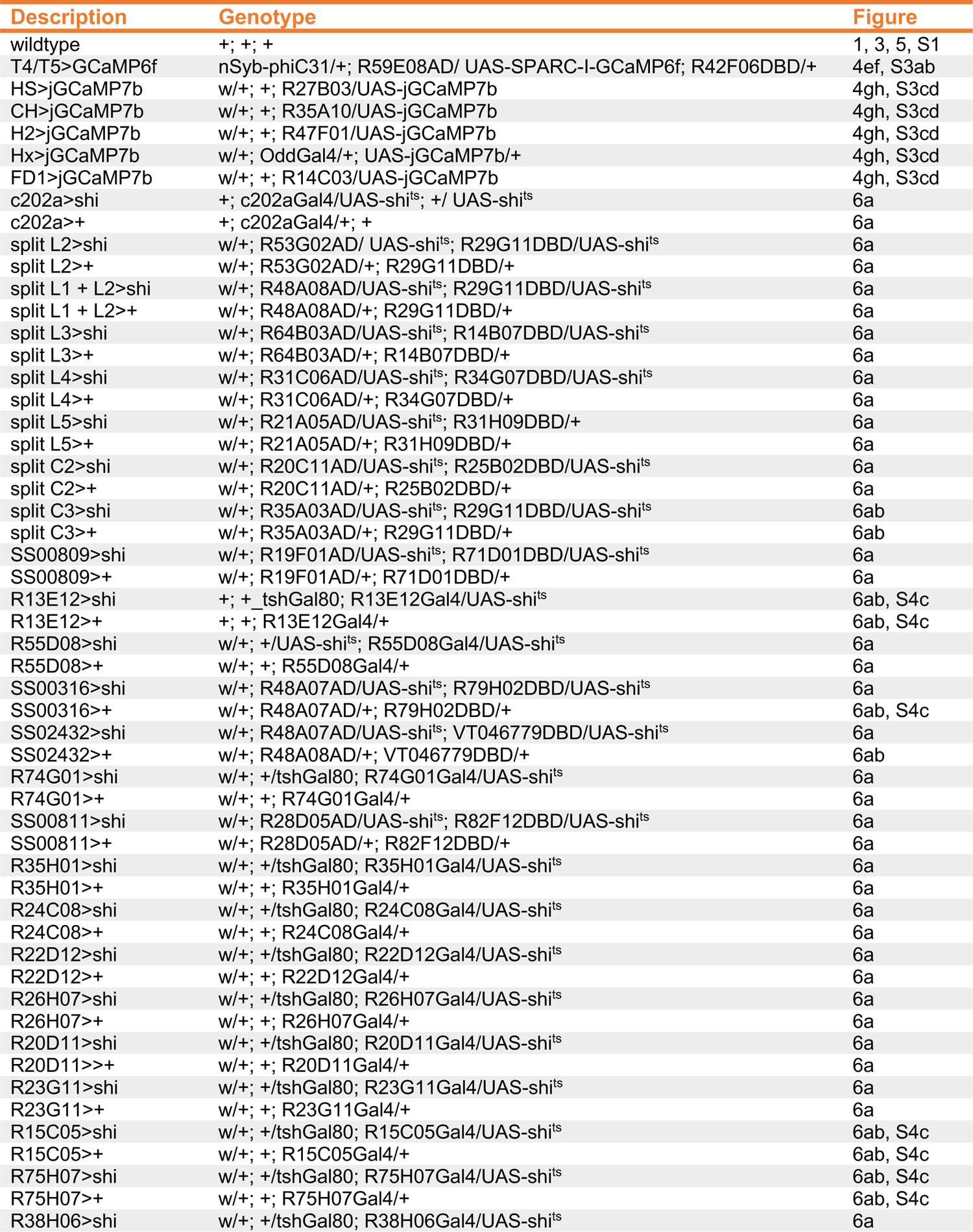
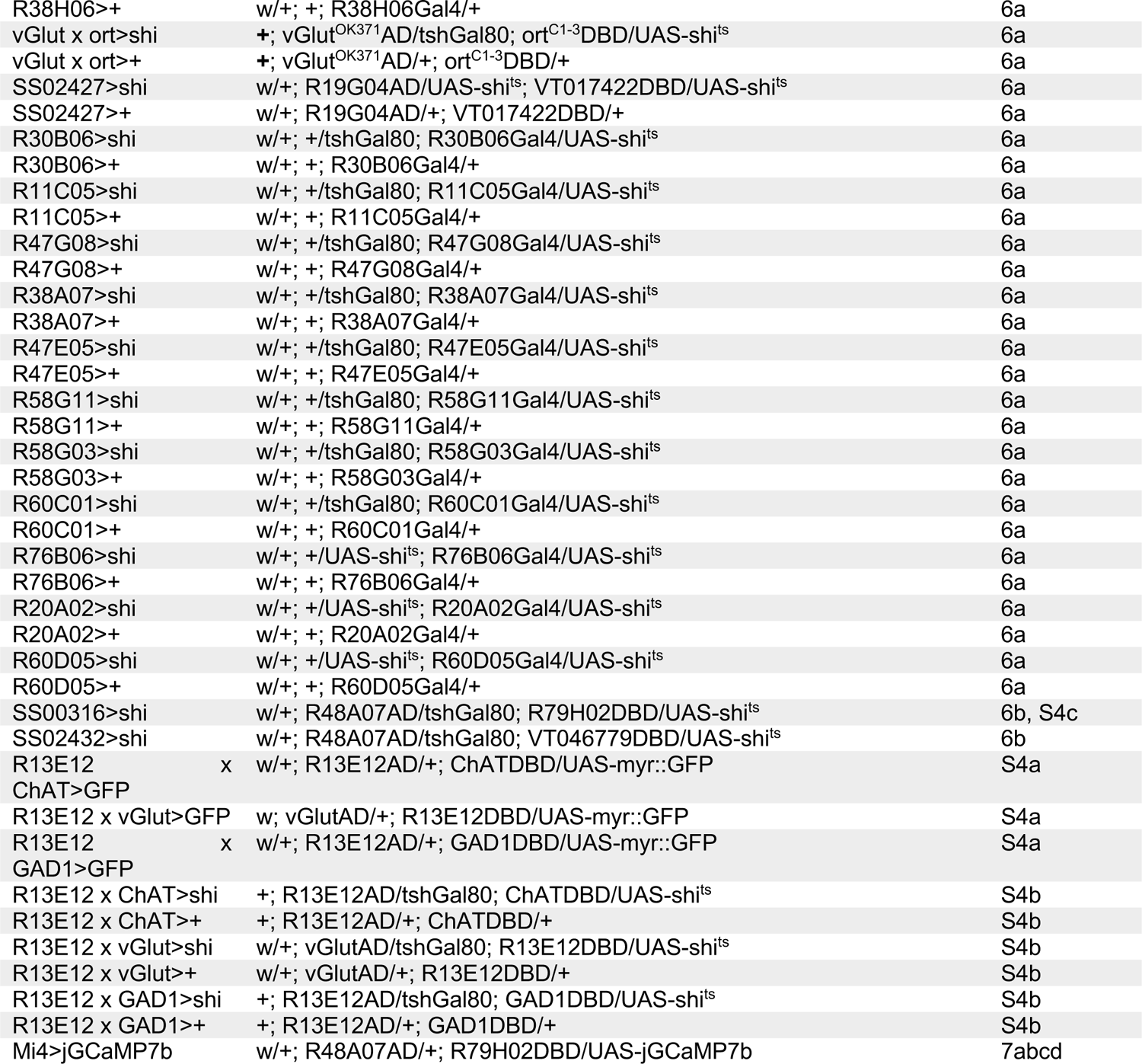
Genotypes of flies used in the experiments.

**Table S2.**
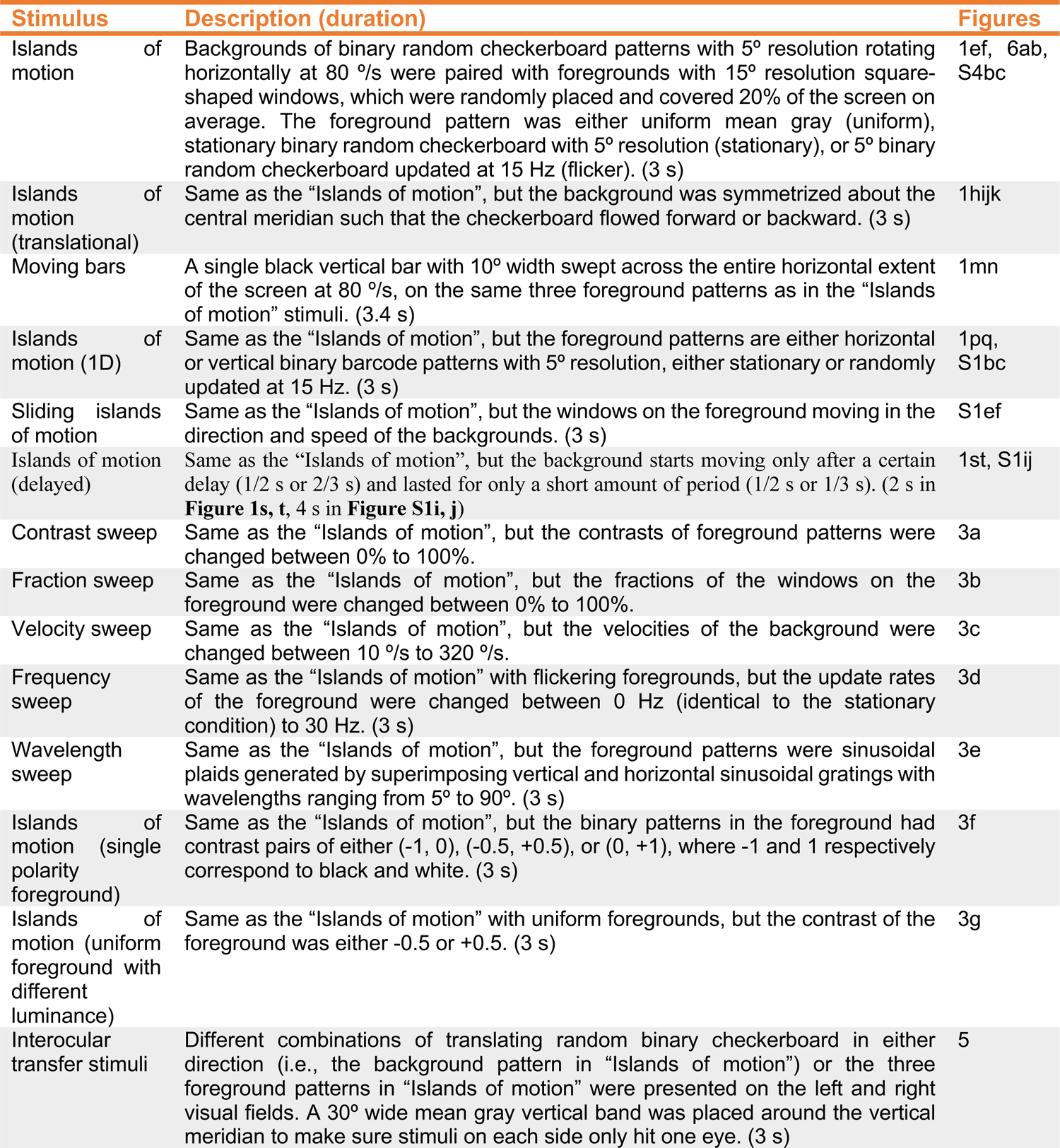
Stimuli for behavioral experiments.

**Table S3.** Stimuli for imaging experiments.

| Stimulus | Description (duration) | Figures |
| --- | --- | --- |
| Localized island of motion | Same as the “Islands of motion” for the behavioral experiment, but the background velocity was 40 °/s and the foreground had a single circular window with 15° diameter. A 5° thick buffer zone with mean gray was set around the window. (2 s) | 4ef, S3ab |
| Islands of motion (imaging) | Same as the “Islands of motion” for the behavioral experiment, but the background velocity was 40 °/s. (2 s) | 4gh |
| Full-field checker | 5° binary random checkerboards, either stationary or randomly updated at 15Hz. (2 or 3 s) | 7abcd, S3c |
| Full-field rotation | 5° binary random checkerboards rotating horizontally at 40 °/s. (2 or 3 s) | 7abcd, S3d |

**Table S4.** Probe stimuli for imaging experiments.

| Stimulus | Description (duration) | Used for |
| --- | --- | --- |
| Island of motion probe | Same as the “Localized island of motion”, but the background moved in the four cardinal directions. (3 s) | T4/T5 |
| Sea of motion probe | Same as “Island of motion probe”, but moving background was presented outside the circular window + the buffer zone, instead of inside. Inside the window remained gray. (3 s) | T4/T5 |
| Local edge probes | A 15° tall, black or white vertical edge moved horizontally at 40 °/s. (3 s) | T4/T5 |
| Square wave probes | Full-contrast vertical square waves with the wavelength of 30° moved horizontally at 30 °/s, interleaved with stationary square waves. (4 s) | HS, CH, H2, Hx, FD1 |
| ON-OFF probe | Full-field, full-contrast, alternating flashes of ON and OFF, each lasting for 3 seconds, repeated for 5 cycles. (6 s) | Mi4 |

